# Xist spatially amplifies SHARP recruitment to balance chromosome-wide silencing and specificity to the X chromosome

**DOI:** 10.1101/2021.10.27.466149

**Authors:** Joanna W. Jachowicz, Mackenzie Strehle, Abhik K. Banerjee, Jasmine Thai, Mario R. Blanco, Mitchell Guttman

## Abstract

Although thousands of lncRNAs are encoded in mammalian genomes, their mechanisms of action are largely uncharacterized because they are often expressed at significantly lower levels than their proposed targets. One such lncRNA is Xist, which mediates chromosome-wide gene silencing on one of the two X chromosomes to achieve gene expression balance between males and females. How a limited number of Xist molecules can mediate robust silencing of a significantly larger number of target genes (∼1 Xist RNA: 10 gene targets) while maintaining specificity to genes on the X within each cell is unknown. Here, we show that Xist drives non-stoichiometric recruitment of the essential silencing protein SHARP (also called Spen) to amplify its abundance across the inactive X, including at regions not directly occupied by Xist. This amplification is achieved through concentration-dependent homotypic assemblies of SHARP on the X and is required for chromosome-wide silencing. We find that expressing Xist at higher levels leads to increased localization at autosomal regions, demonstrating that low levels of Xist are critical for ensuring its specificity to the X chromosome. We show that Xist (through SHARP) acts to suppress production of its own RNA which may act to constrain overall RNA levels and restrict its ability to spread beyond the X. Together, our results demonstrate a spatial amplification mechanism that allows Xist to achieve two essential but countervailing regulatory objectives: chromosome-wide gene silencing and specificity to the X. Our results suggest that this spatial amplification mechanism may be a more general mechanism by which other low abundance lncRNAs can balance specificity to, and robust control of, their regulatory targets.

## INTRODUCTION

In recent years, thousands of lncRNAs have been identified and many have been proposed to regulate gene expression [1–5]. However, their precise mechanisms of action remain largely uncharacterized. One of the key issues is that lncRNAs are generally expressed at low levels such that the number of RNA molecules is less than the number of targets that they are proposed to regulate (sub-stoichiometric) [6–8]. How an individual lncRNA molecule can control multiple distinct targets when it cannot engage with all of them simultaneously remains unknown and has led some to suggest that these lowly expressed lncRNAs may not be functionally important [9,10].

One example of a lncRNA that is expressed at sub-stoichiometric levels relative to its targets is Xist. Expression of Xist is sufficient to induce transcriptional silencing of more than a thousand genes across the >167 million bases of DNA on the X chromosome in order to achieve dosage balance of expression between males and females [11–17]. Previous studies have shown that there are ∼60-200 Xist molecules within an individual cell [18–20], corresponding to an average of ∼1 Xist RNA for every ∼10 genes encoded on the X.

Xist represents an ideal system in which to explore how sub-stoichiometric levels of a lncRNA can regulate its more abundant targets because it is functionally important (developmentally essential) [11,21] with a clear phenotype (transcriptional silencing) [22–24] that occurs at precise and well-defined regulatory targets (X chromosome genes) [15–17]. Recent studies have begun to elucidate the mechanisms by which Xist localizes across the X chromosome and recruits silencing proteins to initiate chromosome-wide silencing. Rather than binding to precise DNA sequences, Xist diffuses from its transcription locus to DNA sites that are in close 3D proximity at both genic and intergenic regions [16,17]. Xist binds directly to SHARP (also called SPEN) [22,25–28], an RNA binding protein that associates with the SMRT and HDAC3 repressive complex to deacetylate histones [29–31], evict RNA Polymerase II [22,24,32], and silence transcription on the X [22,24,25,32–34].

Although these discoveries have uncovered several long-sought molecular mechanisms underlying Xist-mediated silencing, they raise critical new questions about how Xist can achieve the essential quantitative features required for dosage balance. Specifically, Xist-mediated silencing needs to be both specific to ensure that only genes on the X (but not autosomes) are silenced, and robust to ensure that each of the several hundred distinct genes across the X are silenced within each individual cell.

Current models, based on ensemble measurements, cannot explain how Xist achieves these two regulatory objectives – specificity to the X and chromosome-wide silencing – within single cells. For example, Xist localization based on 3D proximity could explain its preferential localization on the X; however, as the X is not partitioned from other chromosomes, this mechanism would not preclude Xist spreading to some autosomal regions within individual cells. Because Xist can silence transcription of genes on autosomes when present in proximity [35–37], its specificity to the X is essential to preclude gene silencing of autosomal genes. Moreover, while Xist localizes broadly across the X when measured in a population of cells [16,17], it cannot localize at all of these positions simultaneously because there is only ∼1 Xist RNA molecule for each megabase of genomic DNA within an individual cell (see Supplemental Note). Accordingly, Xist must localize heterogeneously at distinct locations within individual cells. Such heterogeneous localization would be expected to lead to heterogenous silencing where different genes are silenced in distinct cells. Yet, X chromosome silencing is not heterogenous (see Supplemental Note) [24,38]. Therefore, the stoichiometric silencing model whereby the Xist-SHARP complex localizes at each gene to silence transcription cannot explain how chromosome-wide silencing occurs within single cells.

Here, we explore the mechanisms of how the Xist lncRNA can achieve chromosome-wide gene silencing while ensuring specificity to the X within each individual cell during initiation of XCI.

## RESULTS

### SHARP enrichment on the Xi increases in a non-stoichiometric manner relative to Xist

To explore how sub-stoichiometric levels of Xist can silence genes across the X, we analyzed the temporal and quantitative relationship between localization of Xist and the essential silencing protein SHARP on the inactive X (Xi). SHARP binds directly to Xist and its enrichment on the Xi is dependent on Xist [20,22,24–27]. We reasoned that if SHARP is recruited to the Xi solely through its ability to directly bind to Xist, then the concentration of SHARP would increase proportionally to the concentration of Xist across time (stoichiometric recruitment). In this case, the rate of Xist and SHARP accumulation on the X would be proportional to each other and their ratio would be constant across time (Fig. 1A).

**Figure 1.**
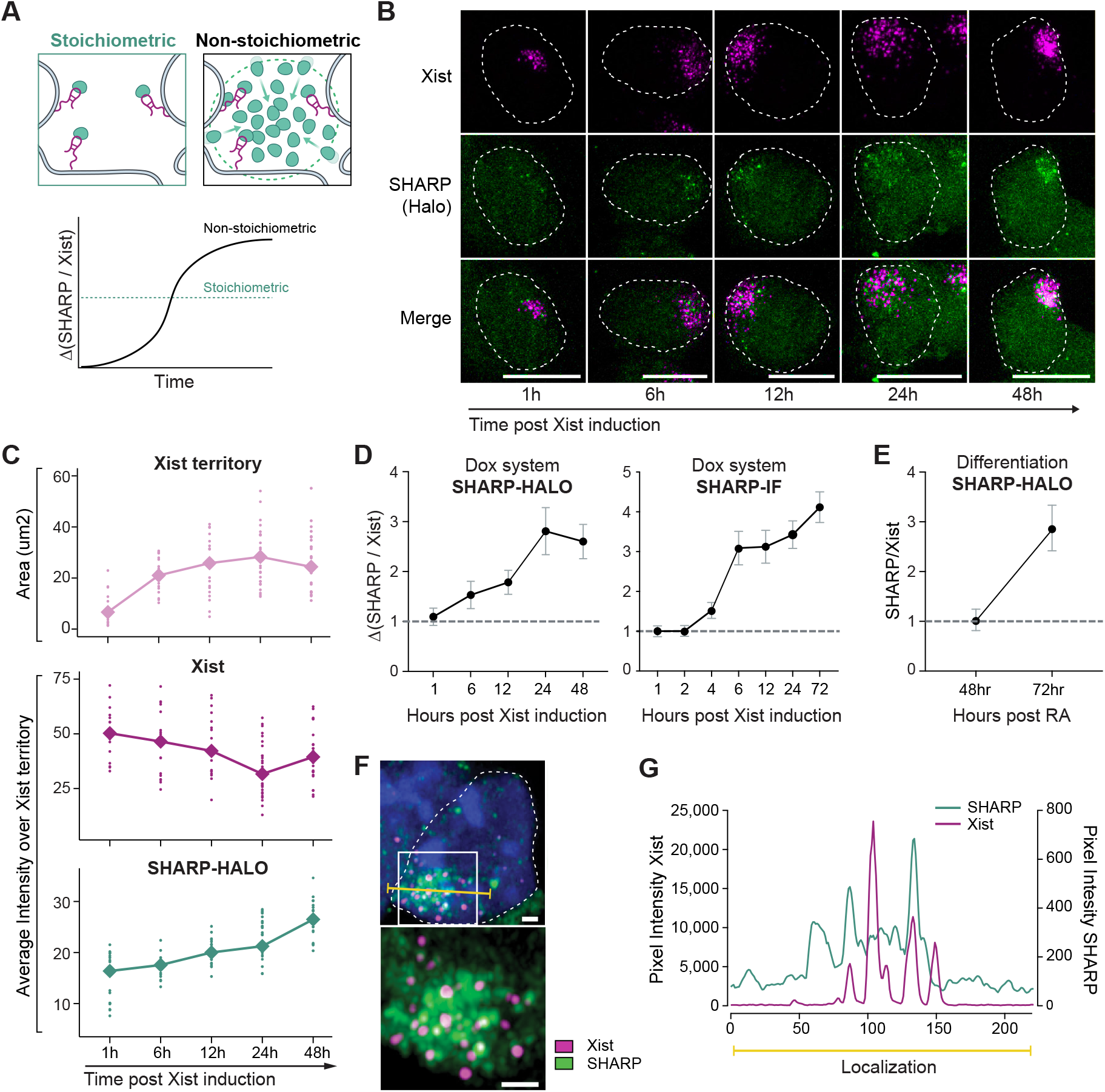
SHARP enrichment over the Xi increases in a non-stoichiometric manner relative to Xist. **(A)** Schematic of two alternative models of Xist-dependent SHARP recruitment to the Xi. Left: stoichiometric, where SHARP (green) localizes through a direct interaction with Xist (magenta); Right: non-stoichiometric, where SHARP can localize even when not directly associated with Xist. Bottom: In the stoichiometric model, the ratio of SHARP to Xist is directly proportional and constant across time; in the non-stoichiometric model, the ratio of SHARP to Xist increases over time in a non-linear manner. **(B)** Representative images of Xist and SHARP localization in TX-SHARP-HALO female mESCs across 48 hours of Xist expression; Xist visualized by RNA-FISH (magenta), SHARP visualized by direct labeling of endogenous SHARP-Halo with Halo ligand (green). Images shown as max. projections; scale bars 10 μm. **(C)** Quantification of temporal dynamics across individual cells (Fig. 1B). Top panel: area of the territory coated by Xist RNA (µm2); Middle panel: average fluorescent intensity of Xist (RNA-FISH) over a unit of Xist territory; Bottom panel: average fluorescent intensity of SHARP (direct Halo labeling) over a unit of Xist territory. Dots represent measurement per individual cell, squares represent mean value for each timepoint. **(D)** Ratio of SHARP to Xist average intensities in TX-SHARP-HALO mESCs at timepoints after Xist induction normalized to one hour group. Left panel: SHARP visualized by direct Halo labeling of endogenous SHARP-Halo across 48 hours; Right panel: SHARP visualized by anti-Halo immunofluorescence of endogenous SHARP-Halo across 72 hours. Dots represent mean per each timepoint, whiskers represent standard deviation. **(E)** Ratio of SHARP (direct Halo labeling) to Xist (RNA-FISH) average intensities upon retinoic acid induced differentiation of TX-SHARP-HALO mESCs normalized to 48 hours group. Dots represent mean per each timepoint, whiskers represent standard deviation. **(F**) Super-resolution imaging of Xist (RNA-FISH; magenta) and endogenous SHARP-Halo (direct Halo labeling; green) in TX-SHARP-HALO mESCs after 24 hours of Xist induction using an Airyscan detector. Top: a single nucleus; Bottom: zoom-in on Xist territory from the top image demarcated by the white box. Images shown as max. projections; scale bars 1 μm. Yellow line shows where the intensity profile (Fig. 1G) was measured. **(G)** Line intensity profile from image in Fig. 1F showing Xist and SHARP intensities across the Xist territory.

To measure this, we used a female F1 hybrid (Bl6 x Cast) mouse embryonic stem cell (mESC) line containing a doxycycline (dox) inducible Xist gene at its endogenous locus and an in-frame HALO tag inserted into both copies of the endogenous SHARP protein (TX-SHARP-HALO cells) [24,39]. This system allows for more temporally synchronized expression of Xist compared to differentiation of female mESCs (Fig. S1A), achieves robust chromosome-wide silencing by 72 hours of induction (Fig. S1B), and utilizes the same molecular components required for initiation of XCI during development and differentiation [23,40]. We induced Xist expression and visualized SHARP (using either a dye that conjugates directly to HALO or an antibody against the HALO tag) along with Xist (using RNA-FISH) across five timepoints (1-48 hours) following dox induction (Fig. 1B; Fig. S1C, S1D). Using both SHARP visualization approaches, we found that the area of the Xist-coated territory increased (Fig. 1C) whereas the total intensity of Xist over the territory increased initially, plateaued, and remained relatively constant (Fig. S1E). This means that the average Xist intensity within the territory decreased over time (Fig. 1C). In contrast, the average intensity of SHARP within the territory continued to increase across all timepoints (Fig. 1C; Fig. S1F; see Methods for quantification). Thus, the ratio of SHARP to Xist intensity is not constant across time, but instead increases in a non-linear manner (Fig. 1D; see Fig.1A for comparison).

To ensure that this effect is not simply a product of our synthetic dox-inducible system (Fig. S1A), we measured the localization of Xist and SHARP across time in female mESCs upon endogenous initiation of XCI using retinoic acid (RA)-induced differentiation. We imaged Xist and SHARP after 2 and 3 days of differentiation (Fig. S1G) and observed a similar relationship between the levels of Xist and the levels of SHARP across time – SHARP levels increased at a faster rate than those of Xist between the two timepoints (Fig. 1E; Fig. S1H). These results demonstrate that SHARP recruitment to the X occurs in a non-stoichiometric manner relative to Xist.

Based on this, we explored whether SHARP is enriched at regions within the Xist-coated territory that are not bound by Xist. To do this, we focused on the Xist territory after 24 hours of dox induction and performed super-resolution imaging of Xist and SHARP (Fig. 1F). We observed foci of Xist within the territory, whereas SHARP exhibits enrichment across the entire territory. As such, there are clear regions of high concentration of SHARP even where Xist is not present (Fig. 1G).

### SHARP forms concentration-dependent assemblies in the nucleus

We next explored how non-stoichiometric recruitment of SHARP to the X might occur. SHARP is an ∼450 kilodalton protein containing four RRM domains [41,42] that bind to Xist [26,32] and a SPOC domain that is critical for recruiting the SMRT and HDAC3 proteins [24,30,31]. The remainder of SHARP is predicted to consist of long intrinsically disordered regions (IDRs; Fig. 2A). Recently, many proteins containing long IDRs have been shown to form concentration-dependent assemblies through multivalent, high-avidity associations [43–46]. Based on this observation, we hypothesized that SHARP might similarly form such concentration-dependent assemblies (Fig. 2B). (Although some concentration-dependent assemblies have been shown to form through phase separation, this is not the only mechanism by which they form [47,48]. In this specific context we are testing whether SHARP forms concentration-dependent assemblies rather than the precise biophysical characteristics of its formation.)

**Figure 2:**
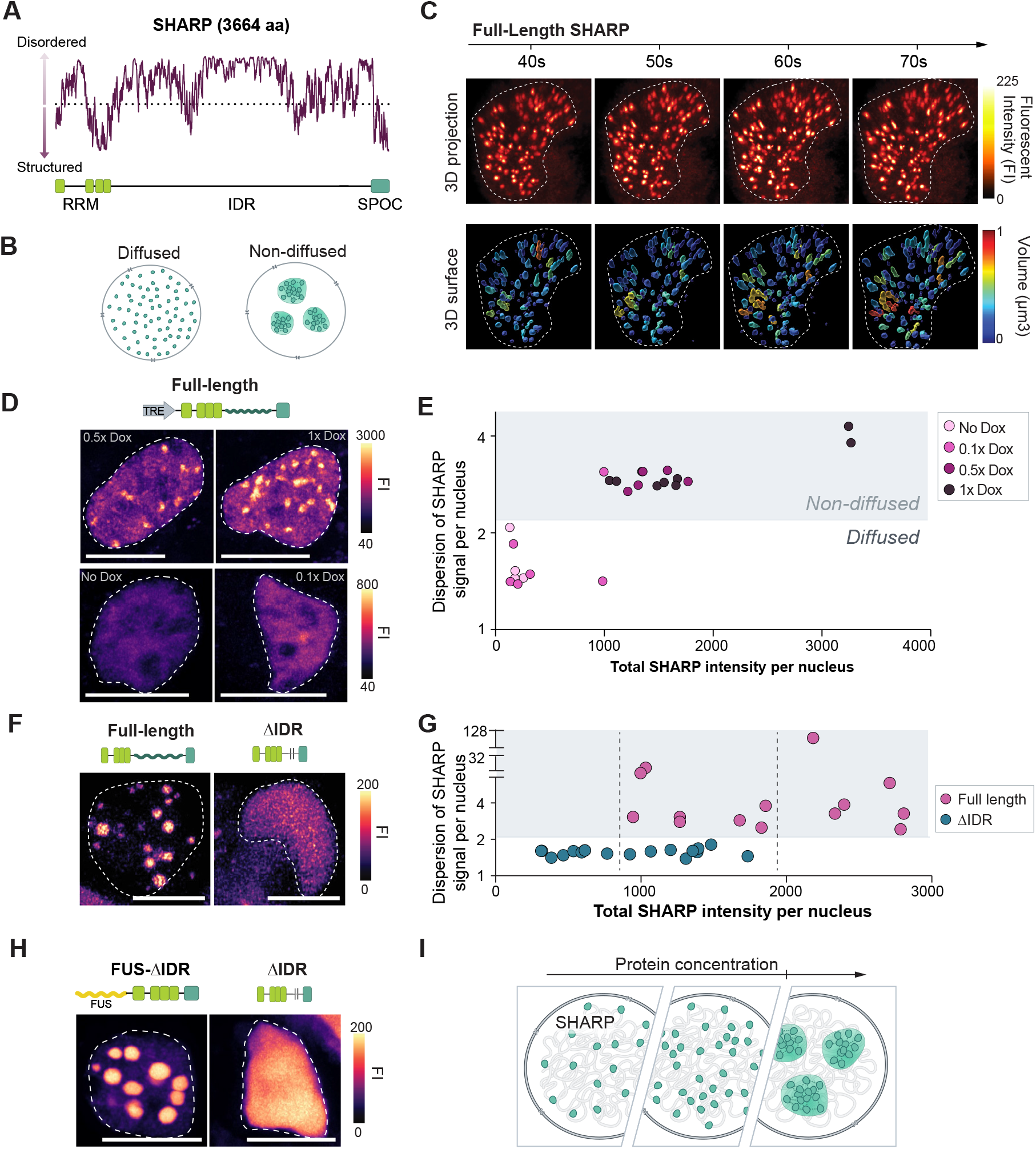
SHARP forms multivalent concentration-dependent assemblies in the nucleus. **(A)** Disordered scores across the SHARP protein using IUPred2 software predictions. Dotted line represents 0.5 probability value for a given structure to be ordered. Bottom visualization demarcates position of known SHARP domains – RNA Recognition motif (RRM; bright green), Spen Paralog and Ortholog C-terminal (SPOC, dark green). **(B)** Schematic representation of molecules within a nucleus organized in a diffused or non-diffused (focal) manner. **(C)** Images across four time-points from a live-cell movie of eGFP-tagged FL-SHARP in transiently transfected HEK293T cells showing non-diffused, focal organization of SHARP molecules. Top panel: 3D reconstructions of the fluorescent intensity signal; Bottom panel: 3D volume reconstructions color-coded based on the size of the condensate; Fluorescent Intensity (FI) **(D)** Images representing localization patterns of eGFP-tagged FL-SHARP in transiently transfected HEK293T cells across increasing expression levels of SHARP (SHARP under dox inducible promoter; dox 1x = 2 µg/mL). Images shown as max. projections; scale bars 10 µm; Fluorescent Intensity (FI). **(E)** Quantification of images (Fig. 1D) plotting the dispersion of SHARP signal across the nucleus versus average SHARP fluorescent intensity per nucleus. **(F**) Representative images of FL-SHARP and ΔIDR-SHARP localization patterns in transiently transfected HEK293T cells. Images shown as max. projections; scale bars 10 µm; Fluorescent Intensity (FI). **(G)** Quantification of images (Fig. 1F) plotting the dispersion of SHARP signal across the nucleus versus average SHARP fluorescent intensity per nucleus. Dashed line represents range of fluorescent intensity that is similar for both groups. **(H)** Images representing localization patterns of mCherry-tagged FUS-ΔIDR-SHARP and eGFP-tagged ΔIDR-SHARP in transiently transfected HEK293T cells. Images shown as max. projections; scale bars 10 µm; Fluorescent Intensity (FI). **(I)** Schematic depicting formation of concentration-dependent SHARP assemblies.

To test this hypothesis, we explored whether SHARP exhibits three known features of multivalent, high-avidity assemblies [43–46]. Specifically, we asked (i) does SHARP form high-concentration foci in the nucleus, (ii) is formation of these foci dependent on the overall concentration of SHARP, (iii) are these foci dependent on multivalent associations mediated through the IDRs, and (iv) are these foci dependent on associations with other molecules of SHARP (homotypic assemblies)?

We expressed full-length SHARP tagged with monomeric eGFP (FL-SHARP; Fig. S2A) in HEK293T cells, a cell type that allows for efficient transfection and controlled expression of the large plasmid containing SHARP and enables characterization of its biochemical and biophysical properties independently of its functional targets. Using this system, we performed live cell imaging and observed that FL-SHARP molecules formed discrete foci within the nucleus (Fig. 2C; Movie S1). These SHARP assemblies also displayed other features of multivalent, IDR-mediated assemblies in that individual molecules exchanged rapidly within a SHARP focus (Fig. S2B) and SHARP foci merged together into larger structures (fusion) or split apart into smaller structures (fission) across time [49,50] (Movie S2; Fig. S2C, S2D).

Next, we used the dox-inducible promoter that drives FL-SHARP expression to titrate its level across a >1,000-fold range of concentrations and determine whether formation of these foci depends on total protein concentration per cell.

We observed that SHARP formed foci only when present at higher concentrations; at low concentrations SHARP was diffuse throughout the nucleus (Fig. 2D, 2E; Fig. S2E; see Methods for quantification) similar to other proteins that do not form assemblies (Fig. S2F).

To determine whether formation of SHARP assemblies is dependent on multivalent interactions driven by its IDRs, we expressed eGFP-tagged SHARP lacking its IDRs (ΔIDR-SHARP; Fig. S2A) in HEK293T cells and imaged its behavior. In contrast to the full-length protein, ΔIDR-SHARP did not form foci (Fig. 2F). Instead, ΔIDR-SHARP localized diffusively throughout the nucleus, even when present at the concentration where FL-SHARP formed foci (Fig. 2G).

Finally, we explored whether these IDR-dependent assemblies form through multivalent associations with other molecules of SHARP (homotypic assemblies) or require sequence-specific associations with other proteins (heterotypic assemblies). To do this, we fused ΔIDR-SHARP to an mCherry-tagged version of the IDR of FUS, an RNA binding protein that is known to form multivalent homotypic associations both *in vitro* and *in vivo* [51–53] (FUS-ΔIDR-SHARP; Fig. S2A) and tested if this synthetic protein rescues the ability of SHARP to form foci independently of its IDRs. We observed that FUS-ΔIDR-SHARP forms assemblies in the nucleus that are comparable to those observed for FL-SHARP (Fig. 2H). While we do not exclude the possibility that the IDRs of SHARP may form heterotypic associations with other molecules, these results indicate that homotypic associations are essential for SHARP to form the observed assemblies.

Together, these results indicate that SHARP forms concentration-dependent assemblies in the nucleus and that formation of these assemblies is dependent on homotypic multivalent interactions driven by its IDRs (Fig. 2I).

### SHARP recruitment to the Xi is dependent on IDR-mediated homotypic associations

To determine if IDR-dependent assemblies of SHARP are critical for its enrichment on the Xi, we tested whether deletion of the IDRs impacts localization over the X.

To do this, we first generated a mESC line containing a deletion of both copies of the endogenous SHARP gene (SHARP-KO; Fig. S3A). In parallel, we utilized mESCs containing an auxin-degradable SHARP (SHARP-AID) [24]. Within each of these lines (SHARP-KO and SHARP-AID), we stably expressed a HALO-tagged or eGFP-tagged version of either full length SHARP (FL-SHARP), SHARP containing a deletion of its RRM domains (ΔRRM-SHARP), or SHARP containing a deletion of its IDRs (ΔIDR-SHARP) (Fig. S3B, S3C). We visualized each of these tagged SHARP proteins along with Ezh2 (to demarcate the Xi) after Xist expression for >72 hours (Fig. 3A; Fig. S3D). As expected, FL-SHARP was enriched over the Xi compartment. In contrast, ΔRRM-SHARP failed to localize on the Xi. Interestingly, we also observed a strong decrease in the enrichment of ΔIDR-SHARP over the Xi, comparable to that observed upon deletion of the RRM domains (Fig. 3B; see Fig. S3E for quantification schematics). The level of Ezh2 was similar in all conditions (Fig. 3B).

**Figure 3:**
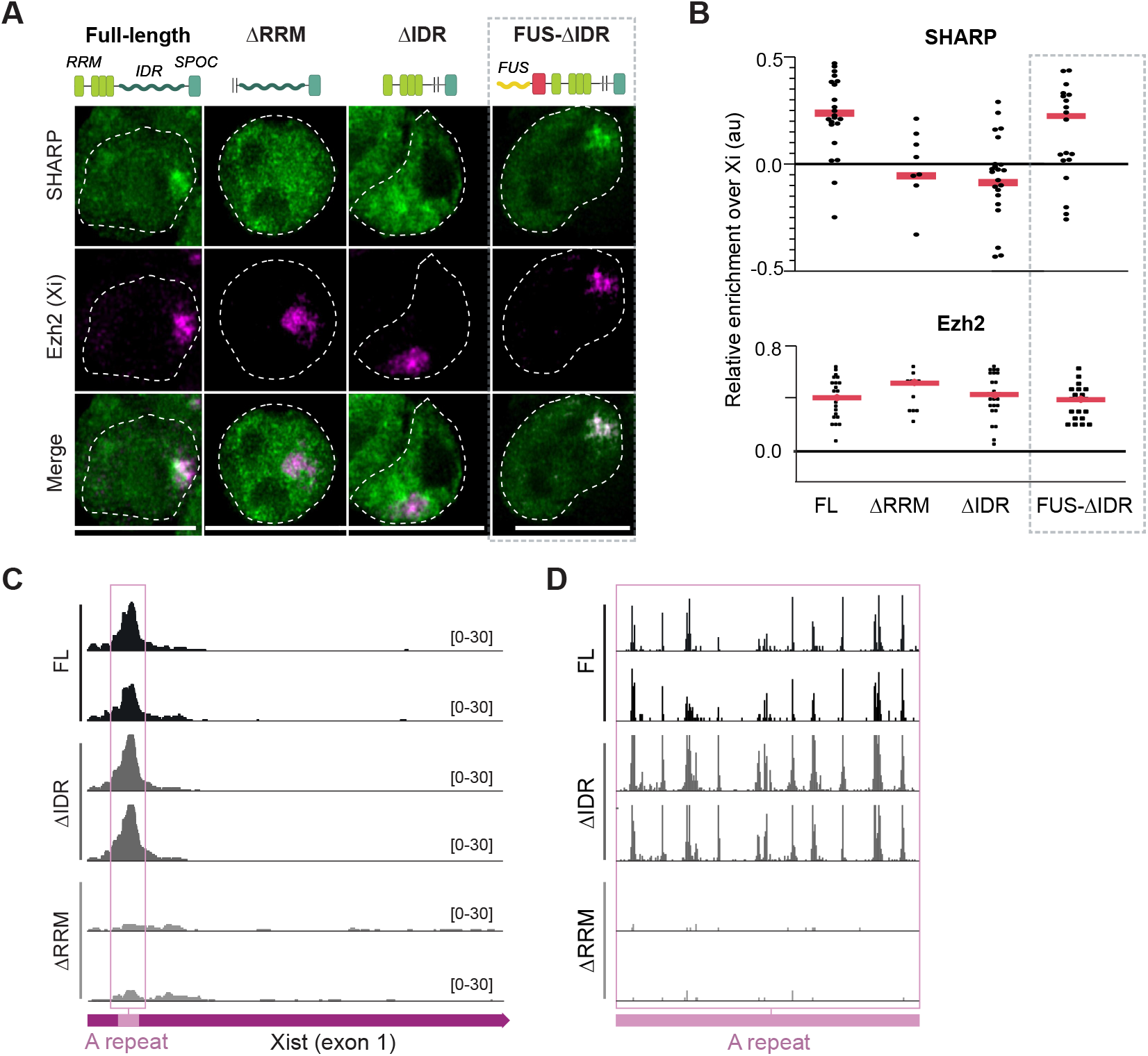
Formation of SHARP assemblies are required for its enrichment on the Xi and dispensable for Xist binding. **(A)** Representative images of SHARP enrichment (eGFP, green) over the Xi (anti-Ezh2 immunofluorescence, magenta) in TX SHARP-KO mESCs containing dox-inducible Xist, genetic deletion of SHARP, and stable integrations of: eGFP-FL-SHARP, eGFP-ΔRRM-SHARP, eGFP-ΔIDR-SHARP, or FUS-mCherry-ΔIDR-SHARP constructs (see Fig. S3B-C for cell lines details). Xist induction for 72h, images shown as z-sections; scale bars show 10 μm. **(B)** Quantification of images (Fig. 3A) plotting (top panel) SHARP fluorescent intensity over the Xi (denoted by Ezh2) normalized to the fluorescent intensity of a random nuclear region of the same size or (bottom panel) Ezh2 fluorescent intensity over the same area normalized to random nuclear region (see Fig. S3D for quantification details). Values for individual nuclei (n>10) are shown; red lines represent median values; 0 represents enrichment not higher than measured over a random nuclear region. **(C)** SHARP enrichment across the first exon of Xist after UV-crosslinking and purification using the HALO tag in female TX SHARP-AID mESCs treated with auxin. Halo-tags were fused to FL-SHARP (top), ΔIDR-SHARP (middle), or ΔRRM-SHARP (bottom, see Fig. S3B-C for cell line details). Two replicates are shown for each cell line; magenta square represents beginning of the first Xist exon, pink square demarcates A-repeat (SHARP binding site). **(D)** Crosslink-induced truncation sites are shown for a zoom-in on the A-repeat region from Fig. 3C demarcated by pink square across three conditions.

Because SHARP binds directly to Xist [22,25–27], we tested whether the ΔRRM- and ΔIDR-SHARP mutants fail to localize on the Xi simply because they cannot bind Xist. To test this, we UV-crosslinked intact cells to form a covalent crosslink between directly interacting proteins and RNA, purified the HALO-tagged SHARP proteins using fully denaturing conditions, and sequenced the associated RNAs (see Methods). We observed that FL-SHARP forms a highly specific interaction with the A-repeat region of Xist. In contrast, expression of ΔRRM-SHARP ablated this interaction across Xist. Interestingly, ΔIDR-SHARP is still able to bind the A-repeat of Xist at comparable levels and positions to that observed for FL-SHARP and the endogenous SHARP protein (Fig. 3C, 3D). These observations are consistent with previous studies that showed that the RRM domains of SHARP are sufficient to bind to Xist [26,32]. Together, these results demonstrate that the IDRs of SHARP are essential for its enrichment on the Xi (Fig. 3A, 3B) even though they are not required for direct binding to Xist (Fig. 3C, 3D).

To exclude the possibility that ΔIDR-SHARP impacts localization on the Xi because it disrupts a cryptic localization domain contained within the protein, we tested whether we could rescue the Xi localization deficits simply by promoting multivalent homotypic associations. To do this, we used our FUS-ΔIDR-SHARP system that forms foci independently of its IDRs (Fig. 2H) and explored whether this could rescue SHARP localization on the X. Indeed, we observed that FUS-ΔIDR-SHARP showed levels of localization over the Xi that were comparable to FL-SHARP after 72 hours of Xist induction (Fig. 3A, 3B). These results demonstrate that the ability of SHARP to form homotypic assemblies (via its IDRs) is essential for its accumulation on the Xi.

### Formation of SHARP assemblies is required for chromosome-wide gene silencing

Because the ΔIDR-SHARP mutant does not accumulate on the Xi, we hypothesized that the ability to form SHARP assemblies is required for Xist-mediated transcriptional silencing during initiation of XCI.

To measure silencing, we performed RNA FISH on Xist and the introns of (i) several genes located across the X that are known to be silenced upon XCI and (ii) genes that are known to escape XCI and therefore remain active upon Xist induction (Fig. 4A; Fig. S4A, Supplemental Note). This single cell readout allows us to restrict our analyses to cells that induce Xist expression (∼50% of cells) and retain both X chromosomes (∼50% of cells; Fig. S4B) [39,54,55]. Of these cells, we found that ∼80% successfully silenced gene expression on one of the two X chromosomes upon Xist induction in wild-type mESCs (Fig. 4B, 4C; Fig. S4C). Next, we measured gene silencing upon genetic deletion (SHARP-KO) or auxin-mediated degradation (SHARP-AID) of SHARP and found that both conditions lead to loss of Xist-mediated transcriptional silencing (Fig. 4B, 4C; Fig. S4C).

**Figure 4:**
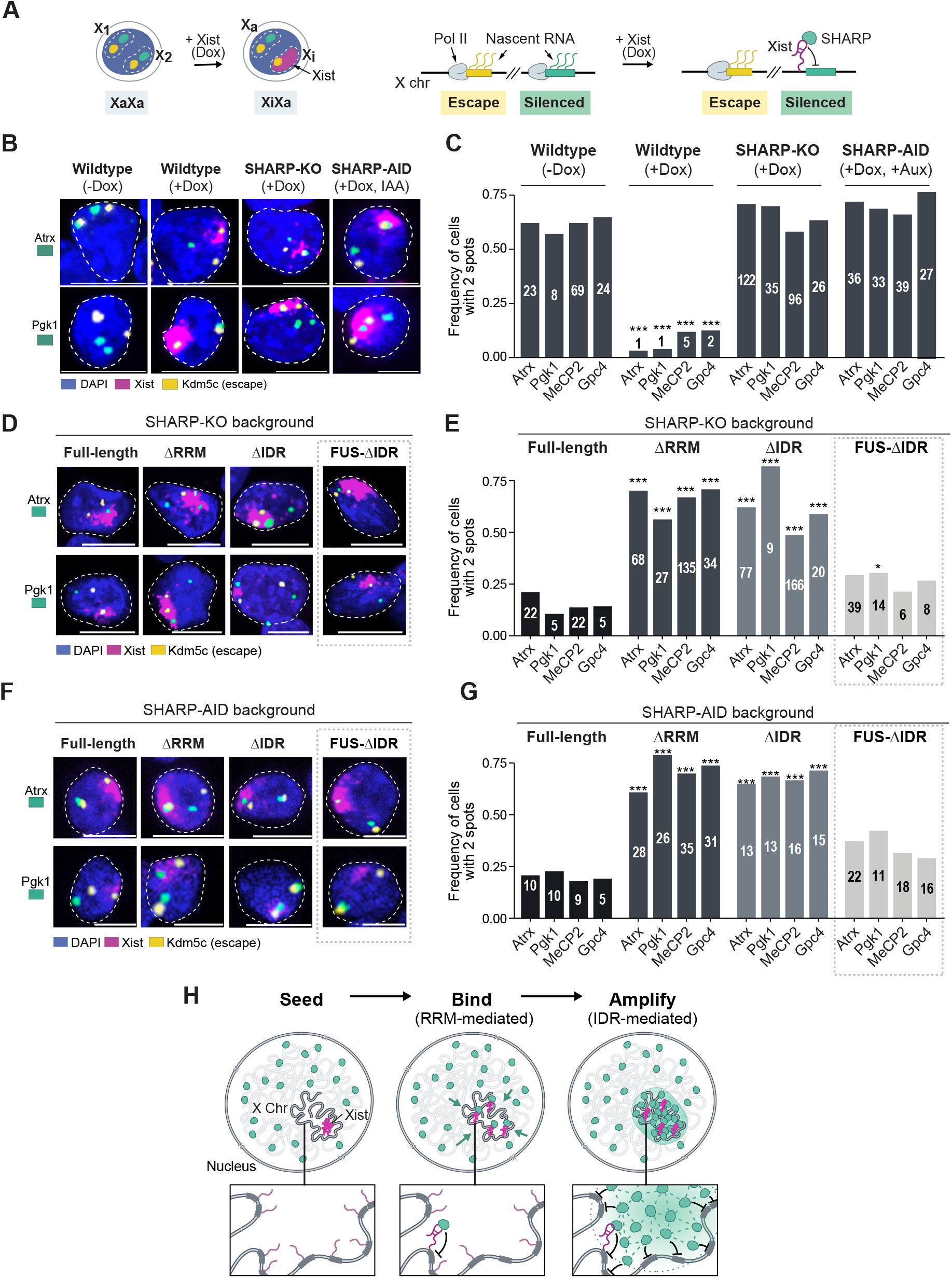
SHARP binding to RNA (via RRM) and formation of assemblies (via IDRs) are both required for chromosome-wide gene silencing. **(A)** Illustration of our RNA FISH measurements in dox-inducible female mESCs. Green: genes that are silenced upon Xist induction; yellow: genes that escape XCI (remain active after Xist induction), magenta: Xist. **(B)** RNA FISH images representing (left to right): wildtype (no dox); wildtype (with dox); SHARP-KO (with dox); auxin-treated SHARP-AID (with dox). Cells were stained for DAPI (blue) and probed for Xist (magenta), escape gene Kdm5c (yellow), and silenced genes Atrx or Pgk1 (green). Images shown as max. projections; scale bars show 10 μm. **(C)** Quantification of multiple RNA FISH images (Fig. 1B) representing the frequency of cells containing two actively transcribed alleles (left to right): wildtype (no dox); wildtype (with dox); SHARP-KO (with dox); auxin-treated SHARP-AID (with dox). **(D**) RNA FISH images for SHARP-KO female mESCs containing stable integrations of (left to right): FL-SHARP, ΔRRM-SHARP, ΔIDR-SHARP, or FUS-ΔIDR-SHARP after >72 hours of dox induction. Cells were stained for DAPI (blue) and probed for Xist (magenta), escape gene Kdm5c (yellow), and silenced genes Atrx or Pgk1 (green). Images shown as max projections; scale bars show 10 μm. **(E)** Quantification of RNA-FISH images representing the frequency of cells containing two actively transcribed alleles for the various SHARP rescue constructs in SHARP-KO female mESCs. **(F)** RNA FISH images for SHARP-AID female mESCs containing stable integrations of (left to right): FL-SHARP, ΔRRM-SHARP, ΔIDR-SHARP, or FUS-ΔIDR-SHARP after >72 hours of dox induction and auxin treatment. Cells were stained for DAPI (blue) and probed for Xist (magenta), escape gene Kdm5c (yellow), and silenced genes Atrx or Pgk1 (green). Images shown as max projections; scale bars show 10 μm. **(G)** Quantification of RNA-FISH images representing the frequency of cells containing two actively transcribed alleles for the various SHARP rescue constructs in SHARP-AID female mESCs. In all panels, only cells containing two escape gene spots (Kdm5c, Kdm6a) and Xist (for dox-induced conditions) were scored for the number of silenced gene spots. Asterisks represent p-values calculated for two proportion Z-test, distributions compared to FL group, * 0.05, ** 0.01, *** 0.001. **(H)** Schematic illustration of the spatial amplification mechanism by which Xist RNA (magenta) can act to amplify SHARP (green) recruitment and gene silencing across the X chromosome.

We measured transcription of the same X-linked genes after stable expression of FL-SHARP, ΔRRM-SHARP, or ΔIDR-SHARP in both SHARP-KO and SHARP-AID backgrounds (Fig. 4D, 4F; Fig. S4D, S4F). As expected, expression of FL-SHARP rescued silencing of these X-linked genes. In contrast, expression of ΔRRM-SHARP failed to silence any of these X-linked genes, consistent with the fact that it can no longer bind to Xist. Importantly, expression of ΔIDR-SHARP also failed to silence these genes (Fig. 4E, 4G; Fig. S4E, S4G). To confirm that silencing depends on the ability of SHARP to form assemblies via its IDRs and not on a specific sequence within the IDRs, we performed the same assay using our SHARP-KO or SHARP-AID cells expressing the synthetic FUS-ΔIDR-SHARP construct that rescues SHARP assembly formation and Xi enrichment (Fig. 2H; Fig. 3A, 3B). We observed a rescue of Xist-mediated transcriptional silencing, comparable to that observed upon expression of FL-SHARP (∼70% silenced cells for FUS-ΔIDR-SHARP versus ∼75% for FL-SHARP) (Fig. 4E, 4G; Fig. S4E, S4G).

Together, these results demonstrate that direct binding of SHARP to Xist (via its RRM domains) and its ability to form concentration-dependent homotypic assemblies (via its IDRs) are both essential and distinct components required for chromosome-wide silencing on the Xi. Our data suggest a spatial amplification mechanism where the direct interaction between Xist (which is enriched on the X chromosome) and SHARP (which is diffusible throughout the nucleus) acts to increase the local concentration of SHARP on the X chromosome. The resulting high local concentrations of SHARP on the X enable formation of IDR-mediated concentration-dependent assemblies that can occur between molecules that are not directly bound to Xist. In this way, these RNA-mediated assemblies can lead to the accumulation of SHARP on the Xi in stoichiometric excess of the number of Xist molecules to enable chromosome-wide silencing (Fig. 4H).

### Xist expression levels are critical for controlling spreading to autosomes

This spatial amplification mechanism explains how Xist can achieve chromosome-wide silencing despite being expressed at sub-stoichiometric levels relative to its target genes (Fig. S5A, S5B, S5C). However, it does not explain why Xist expression levels are low and whether this might be critical for its functional role during XCI. Because Xist spreads to sites on the X based on 3D diffusion from its transcription locus [16,17], we hypothesized that its expression level might control how far it spreads in the nucleus. If true, we would expect that expressing Xist at increasing concentrations would lead to increasing localization of Xist to autosomal regions.

To test this, we used our dox-inducible Xist system, which enables induction of Xist across a range of expression levels by titrating the concentration of dox (Fig. 5A). We induced Xist expression across a range of dox concentrations (referred to as a 0.05x-3x Dox, where 1x = 2 µg/mL), imaged Xist localization in individual cells (Fig. 5B) and observed a strong correlation between Xist expression levels and the area of the nucleus it occupies within individual cells (r=0.75; Fig. 5C). Specifically, Xist occupies on average ∼6.5% of the area of the nucleus when expressed upon RA-induced differentiation (endogenous control). However, cells treated with 3x dox express on average ∼3.4-fold higher levels of Xist (relative to average endogenous levels) and Xist occupies on average ∼23% of the area of the entire nucleus (Fig. 5C).

**Figure 5:**
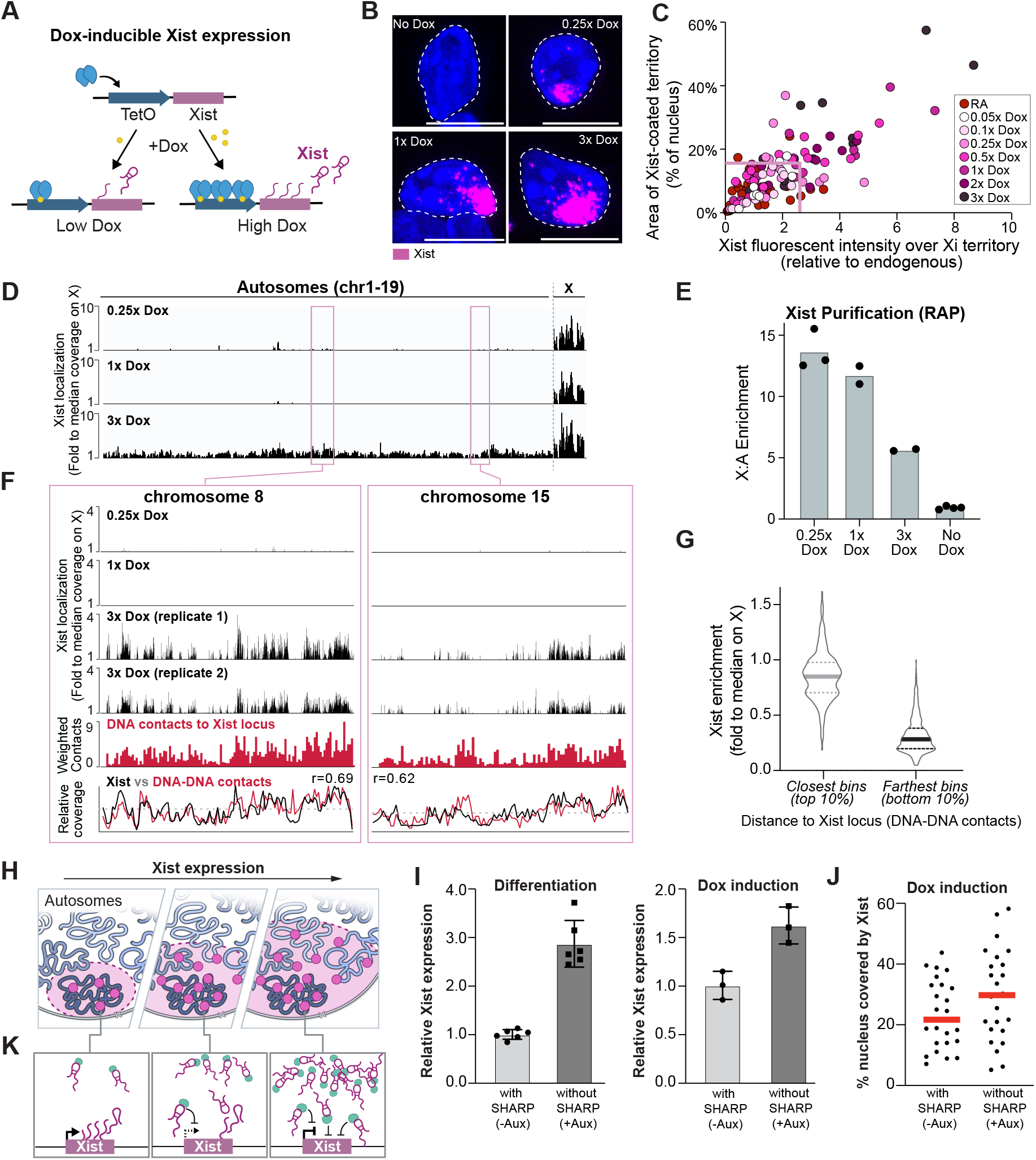
Xist expression levels limit its ability to spread to autosomes. **(A)** Schematic of dox concentration relative to Xist transcription levels in our inducible system. **(B)** Representative images of Xist localization (magenta) within the nucleus (DAPI) of female mESCs treated with increasing concentrations of dox (no dox, 0.25x, 1x, 3x dox). **(C)** Quantification of the area covered by the Xist territory as percent of nucleus (based on RNA FISH and DAPI staining, respectively) relative to Xist expression levels (based on total intensity of Xist per cell) in RA differentiated and dox-induced female mESCs. X-axis: Xist expression levels are normalized to the median expression level of RA differentiated cells; box represents the 95th percentile of Xist expression and area covered in RA differentiated cells. **(D)** Xist enrichment across the genome measured by RNA Antisense Purification (RAP) of Xist followed by DNA sequencing. Scales represent Xist enrichments relative to total genomic DNA in female mESCs treated with 0.25x, 1x and 3x dox. Xist enrichment across the genome normalized to the median coverage across the X (50% of X chromosome bins are 1) *(E*) Overall Xist enrichment on the X chromosome relative to autosomes in 0.25x, 1x, 3x dox-induced and uninduced female mESCs (no dox) as measured by the proportion of sequencing reads that align to the X relative to autosomes (A) in RAP-DNA samples normalized to the expected X:A ratio observed in the unselected genomic DNA sample (input). Dots represent individual RAP-DNA replicates, bars represent mean value. **(F)** Xist enrichment over chr8 (right) and chr15 (left) measured by RAP; Xist enrichment relative to the median coverage on the X (black bars), DNA contact frequency [56] of each region relative to the Xist locus (red bars), bottom: overlay between Xist enrichment and 3D contacts with the Xist locus. **(G)** Levels of Xist enrichment in the 3x dox sample over 1Mb autosomal regions that are closest to Xist locus (left, top 10%) or furthest from Xist locus (right, bottom 10%) based on 3D distance maps calculated from DNA-SPRITE data [56]; Bold lines represent median values, dotted lines represent 25th and 75th percentile. **(H)** Schematic depicting increased Xist spreading in the nucleus with increasing Xist expression levels. **(I)** Relative Xist expression in RA differentiated (left) and dox-induced (right) female SHARP-AID mES cells in the absence or presence of auxin as measured by RT-qPCR. Fold change values were calculated by normalizing to the median of RA differentiated or dox-induced cells in the absence of auxin. Individual points show each replicate value. **(J)** Quantification of percent of nucleus occupied by Xist in dox-induced SHARP-AID cells in the absence or presence of SHARP as measured by RNA FISH (Xist) and DAPI staining (nucleus). **(K)** Model illustrating how Xist (through SHARP) may suppress production of its own RNA at its gene locus through negative feedback.

To determine whether the larger nuclear volumes occupied by Xist correspond to increased localization on autosomes, we performed RNA Antisense Purification (RAP) [16] on Xist and sequenced its associated DNA regions across three different induction conditions (0.25x, 1x, and 3x dox) as well as a negative (no dox) control (Fig. 5D). Because RAP is a bulk measurement, we first confirmed that Xist expression increases in the presence of increasing dox concentrations within a population of cells (using RT-qPCR; Fig. S5D). Then, for each RAP sample, we computed the level of Xist RNA enrichment on the X chromosome (X) by quantifying the proportion of sequencing reads that align to the X relative to autosomes (A). We compared this to the expected X:A ratio observed in the unselected genomic DNA sample (input) (see Methods). In all cases we observed clear enrichment of Xist on the X chromosome (Fig. 5D). However, we observed a steady decrease in X:A enrichment as Xist concentration increased. For example, in samples treated with 3x dox (∼5 fold above endogenous levels) we observed a >2-fold reduction in the X:A enrichment compared to samples treated with 0.25x dox (which approximates endogenous Xist levels) (Fig. 5E).

Because Xist spreads via 3D diffusion, we hypothesized that the autosomal regions that become occupied at increasing dox concentrations are those that are closest to the Xist locus in 3D space. To test this, we computed the 3D contact frequency between the Xist genomic locus and all 1 megabase genomic regions across autosomes [56] (Fig. 5F, S5E). Interestingly, we observed a strong correlation between autosomal regions that are closest to the Xist locus and those that display increased Xist RNA occupancy in the 3x dox condition (Fig. 5G; Fig. S5F). Taken together, these results indicate that sub-stoichiometric expression of Xist (low number of Xist molecules) is a critical mechanism by which cells limit Xist spreading to autosomal regions and ensure its specificity to the X chromosome (Fig. 5H).

Given that low Xist expression levels are critical for ensuring specificity to the X chromosome, we considered possible mechanisms that may act to limit its expression level *in vivo*. One long puzzling observation is that even though Xist and SHARP accumulate in proximity to the Xist transcriptional locus [16,17,24], the Xist gene remains actively transcribed – an essential requirement for XCI. We hypothesized that Xist-SHARP accumulation at its own locus might act to control the level of Xist expression. To test this, we used the auxin degradable SHARP system (SHARP-AID) and measured Xist expression levels upon dox-induction or RA-differentiation. In both cases, we found that depletion of SHARP leads to an ∼2-fold average upregulation of Xist expression (Fig. 5I, Fig. S5G). Consistent with the fact that increased Xist expression leads to an increase in Xist spreading within the nucleus, we observed that degradation of SHARP led to a higher percentage of the nucleus being occupied by Xist (Fig. 5J). Because negative feedback loops often act as regulatory mechanisms to restrict production levels within a defined range [57–60], our results suggest that Xist may act to suppress its own production to ensure specificity to the X (Fig. 5K).

## DISCUSSION

Our results demonstrate a critical spatial amplification mechanism by which Xist balances two essential but countervailing regulatory objectives: specificity to the X and chromosome-wide gene silencing (Fig. 6). We showed that low Xist RNA levels are necessary to ensure specificity to its target sites on the X. Yet, it creates another challenge in that the RNA is expressed at sub-stoichiometric levels compared to its regulatory targets and therefore cannot localize at each of them. We showed that Xist overcomes this challenge by driving non-stoichiometric recruitment of SHARP to amplify its abundance across the X chromosome and enable chromosome-wide gene silencing. Although a stoichiometric model (where Xist recruits SHARP through direct binding and localizes at each of its target genes) would also enable chromosome-wide silencing, it would require Xist to be expressed at dramatically higher levels and therefore reduce Xist specificity to the X. While the spatial amplification mechanism can achieve both specificity and robust silencing, balancing these two competing objectives requires precise quantitative control of Xist RNA levels. Our results highlight a negative feedback loop whereby Xist (through SHARP) may act to suppress production of its own RNA in order to restrict its ability to spread beyond the X (Fig. 6).

**Figure 6:**
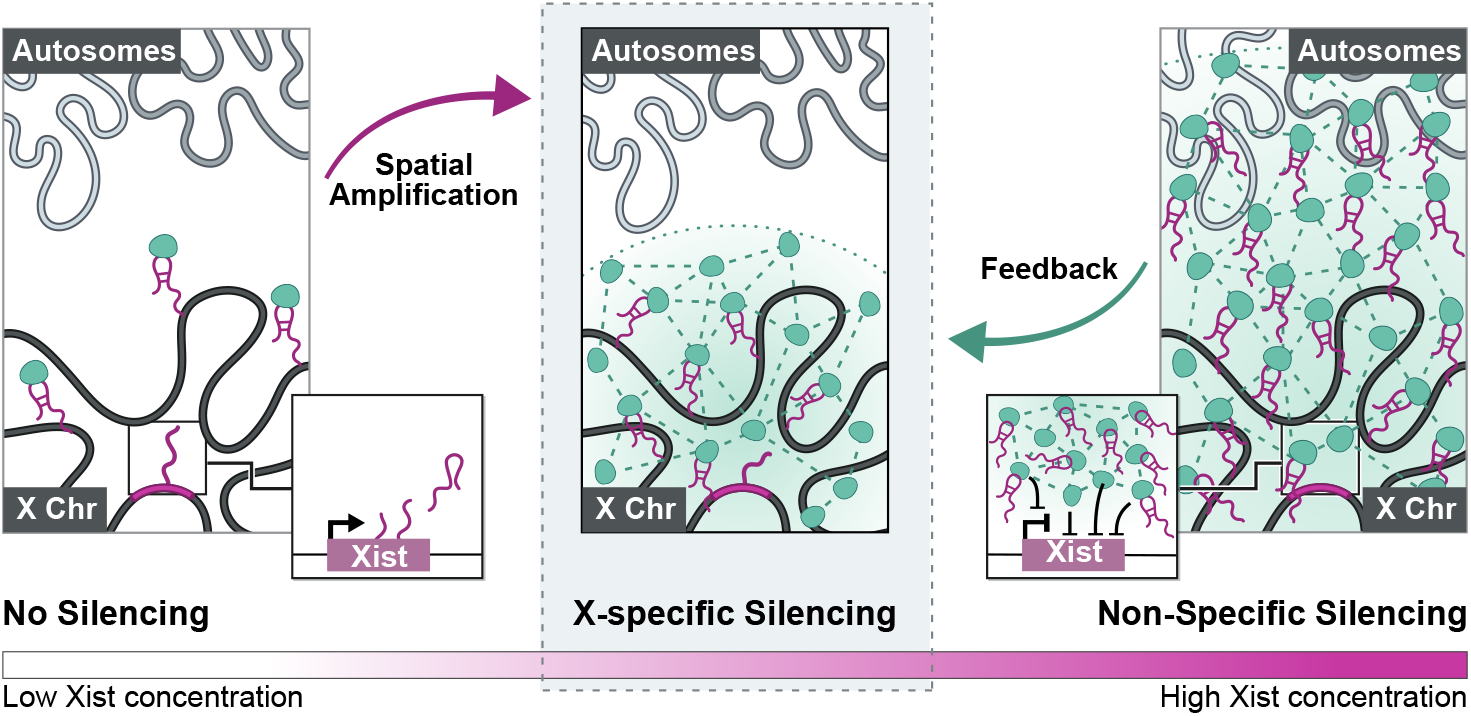
The spatial amplification mechanism balances chromosome-wide silencing and specificity to the X chromosome. A schematic illustrating components of the spatial amplification mechanism. Left: Xist is expressed, accumulates on the X chromosome through 3D diffusion from its transcription locus, binds directly to SHARP and recruits it to the X chromosome in a stoichiometric manner. Zoom-in: At low overall expression, the Xist gene remains actively transcribed. Middle: Once SHARP molecules achieve sufficiently high spatial concentration over X, they form concentration-dependent assemblies (spatial amplification) that enable non-stoichiometric accumulation of SHARP on the X and chromosome-wide silencing. Right: If Xist expression levels get too high, the RNA would start to spread to autosomal regions. At high concentrations, Xist recruits more SHARP molecules to its own locus which can suppress transcription of its own gene (right zoom-panel). This acts to reduce Xist spreading and restrain Xist-SHARP complex on the X chromosome (feedback).

This spatial amplification mechanism is dependent on the fact that Xist can form a high concentration territory on the X through 3D diffusion from its transcription locus (seed). In this way, Xist binding to SHARP increases its concentration on the X (recruit) and enables formation of concentration-dependent protein assemblies that amplify recruitment of repressive proteins to enable chromosome-wide gene silencing (amplify, Fig. 4H). Furthermore, because Xist spreads to its targets via 3D diffusion from its transcription locus, localization specificity is sensitive to its overall expression levels (restriction, Fig. 5H).

Beyond Xist, this spatial amplification mechanism is likely to represent a more general mechanism by which lncRNAs can balance specificity to, and robust control, of their regulatory targets because many lncRNAs share these same properties. Specifically, many hundreds of lncRNAs have been shown to form high-concentration territories in spatial proximity to their transcription sites (seed) and can directly bind and recruit different regulatory proteins (recruit), including those that contain long IDRs [61] (e.g. HP1 [62,63] and SHARP). In this way, lncRNA-mediated recruitment may enable spatial amplification of regulatory proteins and robust regulation of their more abundant targets (amplification). Because many lncRNAs localize in 3D proximity to their targets, low expression levels may similarly be important for ensuring specificity to their genomic DNA targets (restriction).

In this way, spatial amplification may provide a mechanistic answer to two long-standing questions in the lncRNA-field: (i) why many lncRNAs are expressed at relatively low levels and (ii) how low abundance lncRNAs can effectively regulate their more abundant targets.

## ACKNOWLEDGEMENTS

We thank Michael Elowitz, Shasha Chong, Isabel Goronzy, and Drew Honson for their critical comments about the manuscript. We would like to thank Amy Pandya-Jones for sharing her experience with SHARP visualization using pre-extraction followed by immunostaining, and Edith Heard’s laboratory for sharing their cell lines and cell culture protocols. We would like to thank Amy Chow for helpful comments and support with the cell culture work at Guttman laboratory, Biological Imaging Facility at Caltech for their help with microscopy, and Flow Cytometry Facility at Caltech for their help with cell sort. We also thank Shawna Hiley for contributions to writing and editing this manuscript, Marie Bao for her editorial assistance, and Inna-Marie Strazhnik for illustrations. This work was supported by the NIH 4DN program (U01 DA040612 and U01 HL130007), NIH Directors’ Transformative Research Award (R01 DA053178), the New York Stem Cell Foundation, and funds from the California Institute of Technology. M.G. is a NYSCF-Robertson Investigator. J.W.J. was supported by a Caltech BBE post-doctoral fellowship. A.K.B. was funded by NHLBI F30-HL136080 and the USC MD/PhD Program.

## AUTHOR NOTE

While we were working on this manuscript, our long-time collaborators were exploring the localization of various proteins involved in XCI using super-resolution microcopy. In parallel, they made a similar observation regarding the dynamics of Xist and SHARP localization and recently reported this in Markaki et al [19]. Some of the authors on this manuscript are also authors on that manuscript because we openly shared several reagents used for their study, including the SHARP mutant constructs that we generated for this manuscript.

## AUTHOR CONTRIBUTIONS

J.W.J. conceived of this project with M.G. J.W.J. and M.S. performed experiments, analyzed and interpreted data, generated figures, and wrote the paper. A.K.B. performed all CLAP sequencing experiments and provided comments and edits for the manuscript. J.T. created the SHARP rescue constructs with A.K.B and assisted with cell culture. M.R.B worked with A.K.B on CLAP sequencing experiments, worked with J.W.J on RAP sequencing experiments, analyzed sequencing data, and provided comments and edits for the manuscript. M.G. conceived of this project with J.W.J. and oversaw all experiments and analysis, performed analysis and generated figures, and wrote the paper with J.W.J. and M.S.

## SUPPLEMENTAL NOTE

### Xist localization across the X and chromosome-wide silencing

Previous studies have shown that there are between 60-200 copies of Xist within each nucleus [18,19]. This level of expression is sufficient to drive chromosome-wide silencing across the >1500 genes encoded on the X. Based on these numbers, Xist cannot simultaneously localize to each gene within each cell because there are not enough Xist molecules present; it must instead mediate silencing over several genes at once (Fig. S5A). As such, Xist localization within individual cells must be heterogenous such that in one cell it localizes at a subset of genes but in another cell it localizes at a different subset of genes.

Based on ensemble measurements, we know that Xist does not preferentially accumulate at specific sequences (e.g., promoters) but instead localizes broadly across the chromosome (Fig. S5B). This means that the Xist RNA molecules within each cell must localize randomly at distinct positions spread across the >167 megabases of the chromosome.

Using this information, we can simulate the expected occupancy of Xist across the X within single cells in a manner that would explain the ensemble pattern (Fig. S5B). We find that the likelihood that Xist is present over any given gene within an individual cell is extremely low (on average <5% of genes per cell would be covered by Xist) (Fig. S5C). For example, Xist would be expected to localize over any region of Pgk1 in only ∼7% of cells (Fig. S5C). As such, if Xist-mediated silencing was solely dependent on such localization, we would expect that this gene would remain active in >90% of individual cells. However, using our single cell measurements we observe that this gene is silenced in >87% of single cells (Fig. S4C). Therefore, these single cell measurements allow us to measure chromosome-wide silencing when focusing on a subset of X chromosome genes.

**Figure S1:**
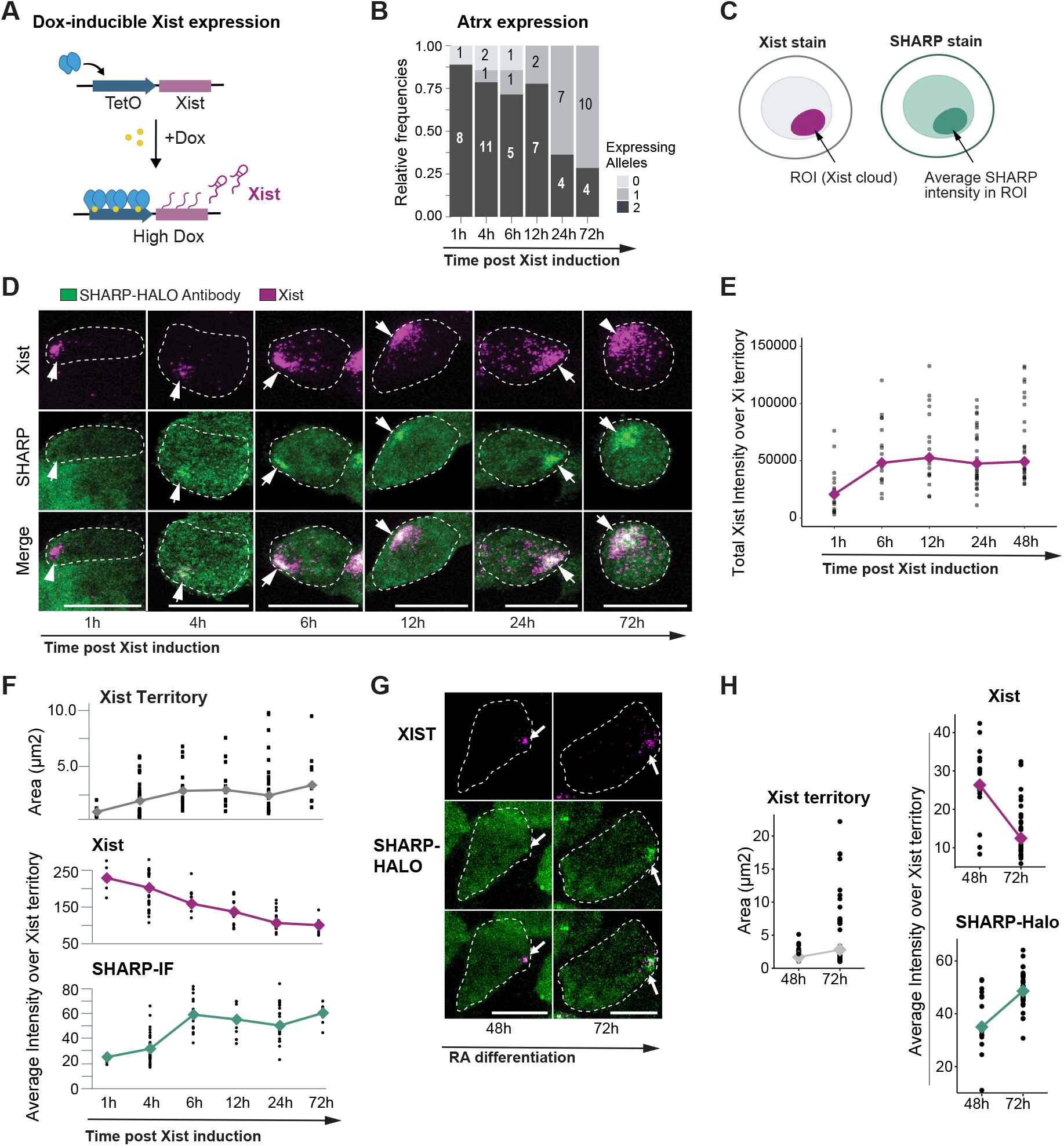
SHARP enrichment over the Xi increases in a non-stoichiometric manner relative to Xist. **(A)** Schematic of dox-inducible Xist expression system. The endogenous Xist promoter is replaced with a TetO element that can be activated upon the addition of doxycycline. **(B)** Percent of cells expressing zero, one, or two alleles of the silenced Atrx gene as measured by RNA-FISH at various timepoints after Xist induction. Only cells containing two escape gene spots (Kdm5c) and Xist were scored. **(C)** Illustration of SHARP enrichment analysis over the Xi in TX-SHARP-HALO female mESCs. The Xi region was demarcated based on Xist RNA-FISH, SHARP was demarcated by either direct HALO labelling or immunofluorescence (anti-HALO). Fluorescent intensities of RNA-FISH probes, HALO tag, or anti-HALO immunofluorescence were then quantified within the defined Xi region and plotted. **(D)** Representative images of Xist and SHARP localization in TX-SHARP-HALO female mESCs across 72 hours of Xist expression; Xist visualized by RNA-FISH (magenta), SHARP visualized by immunofluorescence labelling with anti-HALO antibody (green). Images shown as max. projections; scale bars show 10 μm. **(E)** Quantification of total fluorescence intensity of Xist (RNA-FISH) in multiple individual cells over 48 hours of Xist expression (Fig. 1B). **(F)** Quantification of Xist and SHARP intensities in multiple individual cells over 72 hours of Xist expression (Fig. S1D). Top panel: area of the territory coated by Xist RNA (µm2); middle panel: average fluorescent intensity of Xist (RNA-FISH) per unit within the Xist territory; bottom panel: average fluorescent intensity of SHARP (anti-Halo antibody) per unit within the Xist territory. **(G)** Representative images of Xist and SHARP localization in TX-SHARP-HALO female mESCs after 48 and 72 hours of RA-induced differentiation; Xist visualized by RNA-FISH (magenta), SHARP visualized by direct Halo labelling (green). Images shown as max. projections; scale bars show 10 μm. **(H)** Quantification of Xist and SHARP in individual differentiated cells (Fig. S1G). Left panel: area of the territory coated by Xist RNA (µm2); top right panel: average fluorescent intensity of Xist (RNA-FISH) per unit within the Xist territory; bottom right panel: average fluorescent intensity of SHARP (direct Halo labelling) per unit within the Xist territory.

**Figure S2:**
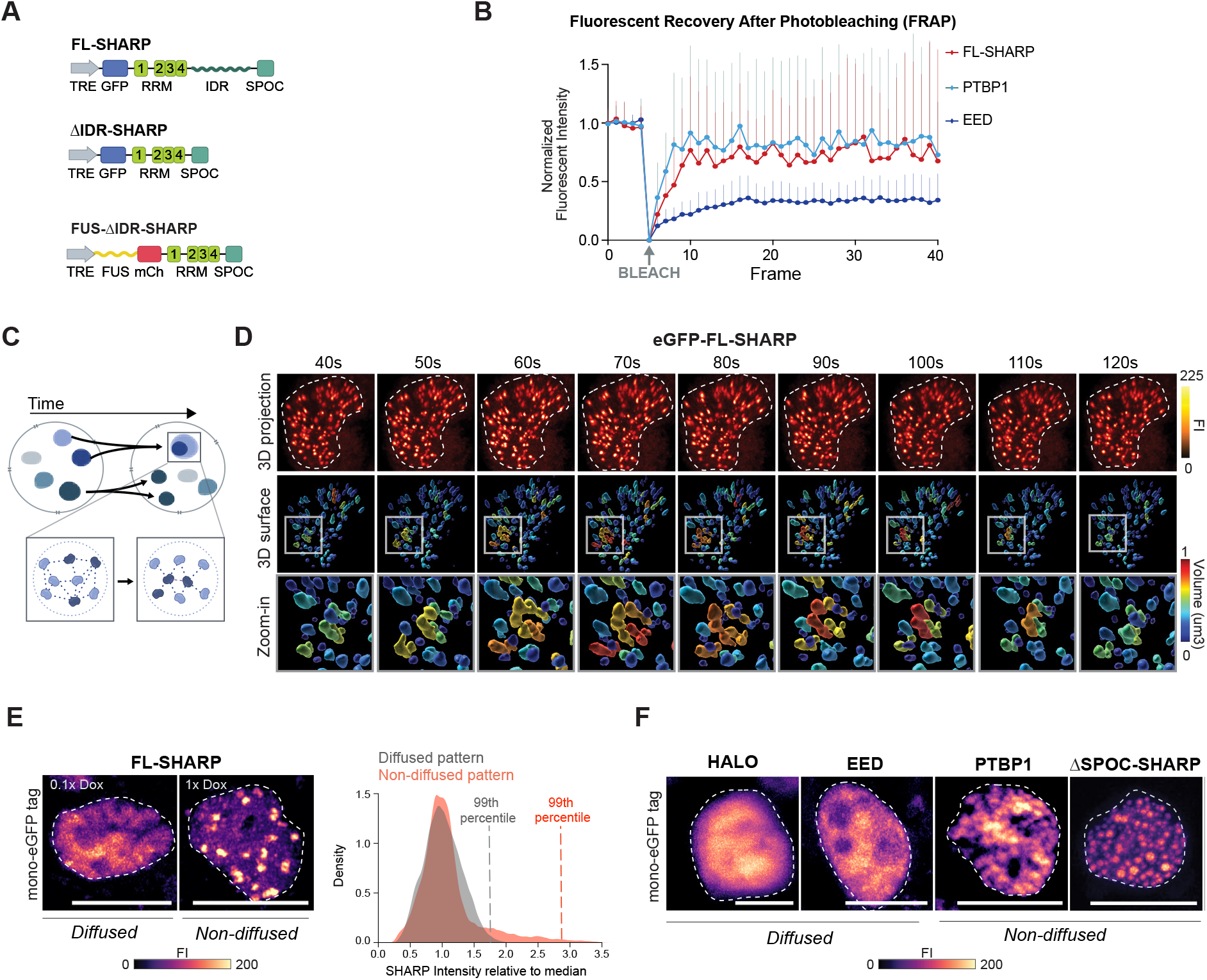
SHARP forms multivalent concentration-dependent assemblies in the nucleus. **(A)** Schematic of the domains included in the eGFP-tagged FL-SHARP and ΔIDR-SHARP, and the mCherry-tagged FUS-ΔIDR-SHARP rescue construct used in Fig. 2 and Fig. S2. **(B)** FRAP recovery curve of eGFP-tagged FL-SHARP (red), positive control PTBP1 (forms assemblies; light blue), and negative control EED (does not form assemblies; dark blue). Error bars represent standard deviation of at least five replicates. **(C)** Schematic depicting physical characteristics of concentration-dependent assemblies, including foci formation, fission and fusion, and rapid diffusion of proteins within an assembly (inset). **(D)** Images across nine time-points from a live-cell movie of eGFP-tagged FL-SHARP in transiently transfected HEK293T cells (Movie 1, Movie 2) showing non-diffused, focal organization of SHARP molecules. Top panel: 3D reconstructions of the fluorescent intensity signal; middle panel: 3D volume reconstructions color-coded based on the volume of the focus; bottom panel: zoom-in representing one region of the nucleus that changes volume; Fluorescent Intensity (FI). **(E)** Comparison of diffused or non-diffused localization patterns of FL-SHARP at different dox concentrations. Left: images representing FL-SHARP expressed with either 0.1x dox (diffused) or 1x dox (non-diffused) in transiently transfected HEK293T cells; images shown as max projections; scale bars show 10 μm.; Right: Histograms representing fluorescent intensities for two cells showing diffused and non-diffused localization patterns. The intensity at the 99th percentile of each distribution is shown with the dashed lines. **(F)** Images representing nuclear localization pattern of eGFP-tagged proteins in transiently transfected HEK293T cells. On the left: proteins that have not been reported to form assemblies (HALO and EED), on the right: an eGFP tagged protein that has been reported to form assemblies (Ptbp1) and ΔSPOC-SHARP that also forms assemblies. Images shown as max projections; scale bars show 10 μm.

**Figure S3:**
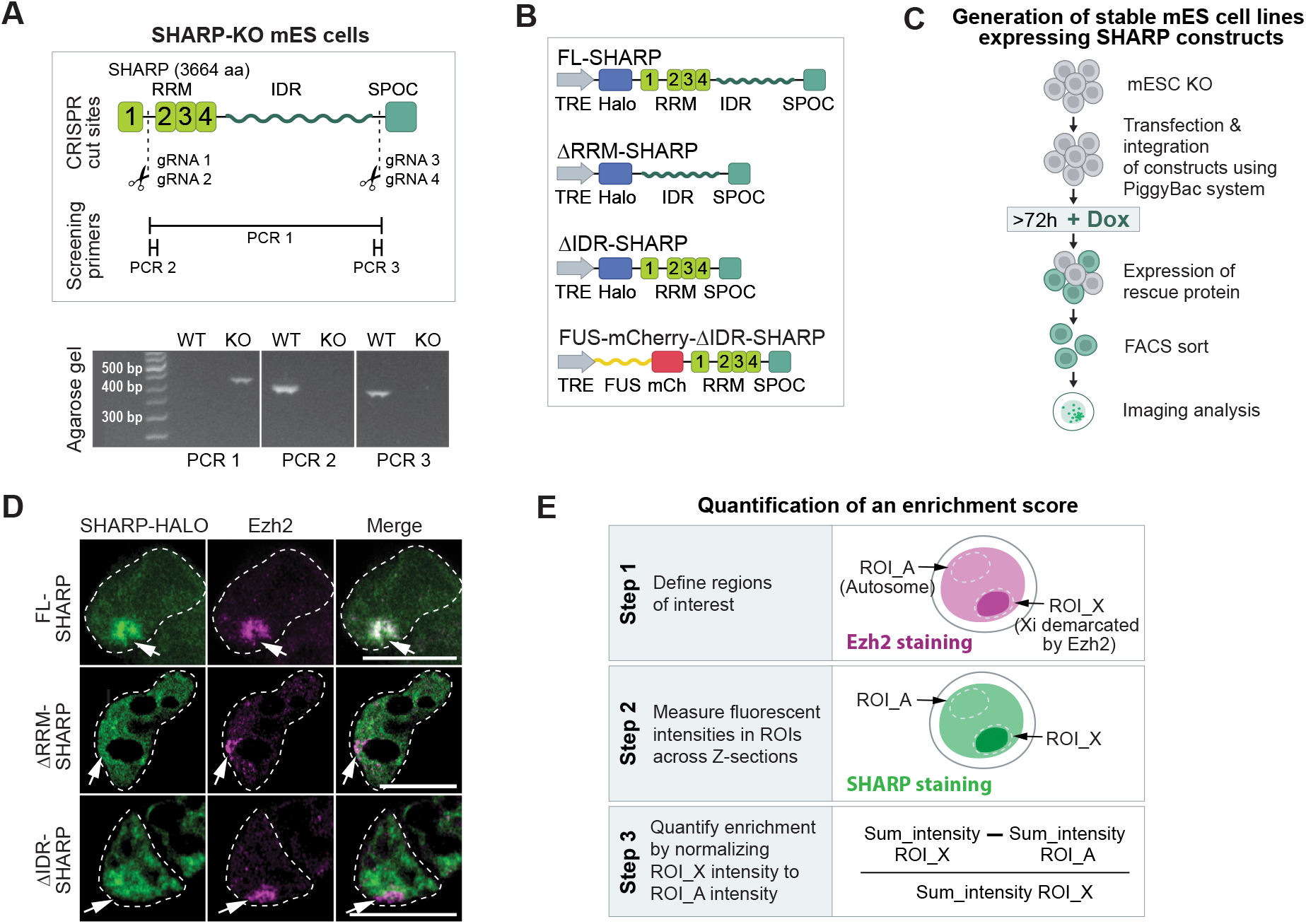
Formation of SHARP assemblies is required for SHARP enrichment on the Xi and is dispensable for Xist binding. **(A)** Generation of SHARP-KO cell line in TX mESCs. Top: schematic of CRISPR cut sites used to generate SHARP-KO mESCs and PCR primers used to screen for KO clones; bottom: agarose gel confirming homozygous deletion of SHARP in SHARP-KO clone H8 mESCs. **(B)** Schematics of constructs used to generate rescue cell lines in TX SHARP-KO or TX SHARP-HALO-AID backgrounds. Grey arrow represents dox-inducible promoter; blue box represents HALO (or eGFP) tags used; light green boxes represent RNA Recognition Motifs (RRM); wavy green line represents the Intrinsically Disordered Regions (IDRs); dark green box represents the Spen Paralog and Ortholog C-terminal (SPOC) domain. Full-length SHARP (FL-SHARP), deletion of RRM domain (ΔRRM-SHARP), deletion of IDR domain (ΔIDR-SHARP), deletion of IDR domain and insertion of alternative IDR domain from FUS protein (FUS-ΔIDR-SHARP). **(C)** Schematic showing experimental workflow for generating and enriching stable SHARP rescue mESCs (FL-SHARP, ΔRRM-SHARP, ΔIDR-SHARP, FUS-ΔIDR-SHARP) using constructs from Fig. S3B. **(D)** Representative images of SHARP enrichment (HALO, green) over the Xi (anti-Ezh2 immunofluorescence, magenta) in female mESCs containing dox-inducible Xist, genetic deletion of SHARP, and stable integrations of HALO-tagged FL-SHARP, ΔRRM-SHARP, or ΔIDR-SHARP. Xist and SHARP rescue constructs induced with doxycycline for 72 hours; images shown as Z-sections; scale bars show 10 μm. **(E)** Diagram of image analysis workflow for quantifying SHARP enrichment over the Xi (Fig. 2B).

**Figure S4:**
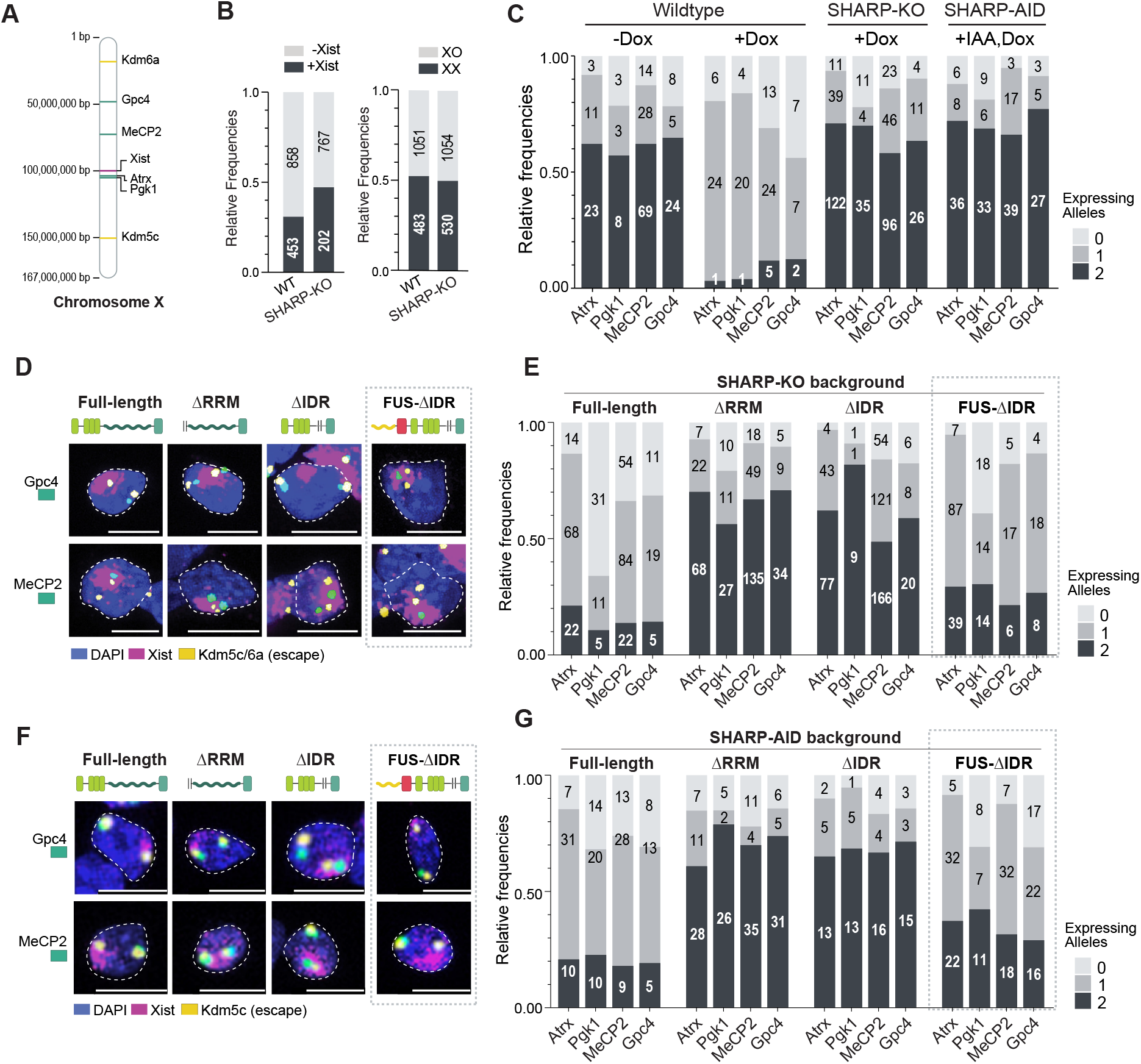
SHARP binding to RNA (via RRM) and formation of assemblies (via IDRs) are both required for chromosome-wide gene silencing. **(A)** Schematic of mouse X chromosome showing the locations of the various genes probed in RNA-FISH experiments. **(B)** Frequency of Xist induction (left) and X chromosome ploidy (right) in wildtype and SHARP-KO mESCs based on quantification of RNA-FISH images. **(C)** Quantification of RNA-FISH images (Fig. 4B) representing the frequency of cells containing two, one, or zero actively transcribed alleles. Left to right: wildtype (-dox); wildtype (+dox); dox-induced SHARP-KO; dox-induced, auxin-treated SHARP-AID female mESCs. Only cells containing two escape gene spots (Kdm5c, Kdm6a) and Xist (for dox-induced conditions) were scored for the number of silenced gene spots. **(D)** RNA-FISH images from SHARP-KO female mESCs containing stable integrations of (left to right): FL-SHARP, ΔRRM-SHARP, ΔIDR-SHARP, or FUS-ΔIDR-SHARP after >72 hours of dox induction. Cells were stained for DAPI (blue) and probed for Xist (magenta), escape gene Kdm5c (yellow), and silenced genes Gpc4 or MeCP2 (green). Images shown as max projections; scale bars show 10 μm. **(E)** Quantification of RNA-FISH images representing the frequency of cells containing two, one, or zero actively transcribed alleles for the various SHARP rescue constructs in SHARP-KO female mESCs. Only cells containing two escape gene spots (Kdm5c, Kdm6a) and Xist (for dox-induced conditions) were scored for the number of silenced gene spots (Fig. S4D). **(F)** RNA-FISH images from SHARP-AID female mESCs containing stable integrations of (left to right): FL-SHARP, ΔRRM-SHARP, ΔIDR-SHARP, or FUS-ΔIDR-SHARP after >72 hours of dox induction. Cells were stained for DAPI (blue) and probed for Xist (magenta), escape gene Kdm5c (yellow), and silenced genes Gpc4 or MeCP2 (green). Images shown as max projections; scale bars show 10 μm. **(G)** Quantification of RNA-FISH images representing the frequency of cells containing two, one, or zero actively transcribed alleles for the various SHARP rescue constructs in SHARP-KO female mESCs. Only cells containing two escape gene spots (Kdm5c, Kdm6a) and Xist (for dox-induced conditions) were scored for the number of silenced gene spots (Fig. S4F).

**Figure S5:**
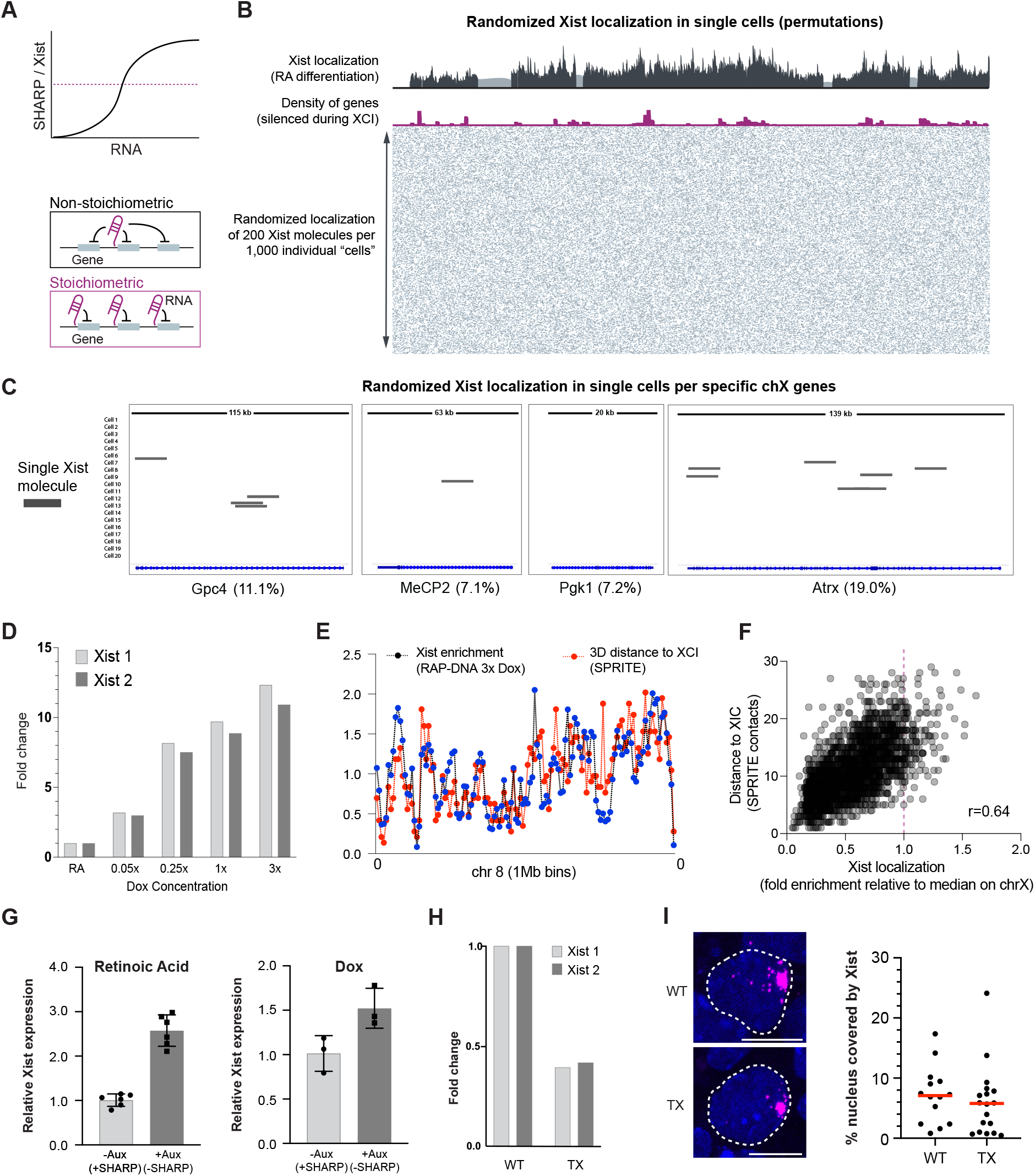
Xist expression levels limit its ability to spread to autosomes. **(A)** Top: schematic depicting expected ratios of SHARP to Xist based on increasing concentration of Xist RNA; bottom: diagrams illustrating non-stoichiometric and stoichiometric gene silencing by Xist. **(B)** Randomized Xist localization in simulated single cells (permutations) compared to experimental data. Top panel: Xist localization after 48 hours of RA-differentiation in female mESCs from bulk RAP-DNA experiments (16); middle panel: gene density across X, only genes that undergo XCI are plotted; bottom panel: randomized localization of 200 Xist molecules in 1000 random “cells”; Xist represented by black squares. **(C)** Simulation visualizing localization of Xist molecules over genes that undergo transcriptional silencing during XCI across 20 cells (zoom-in from Fig. S5B). Percent reflects proportion of Xist molecules overlapping the gene in all 1000 permutations. **(D)** Expression levels of Xist in several populations of cells treated with increasing dox concentrations as measured by RT-qPCR using two different primer pairs (Xist 1 and Xist 2). Samples normalized to i) RA-differentiated levels and ii) GAPDH levels. **(E)** Comparison of Xist occupancy (black lines; 3x dox RAP data) and DNA contact frequency with Xist locus (red lines, SPRITE data) across 1Mb DNA regions of chr8. Xist enrichment and 3D distance are normalized to their median coverage across chromosome 8 to place them on the same relative scale. **(F)** Scatterplot representing the frequency of 3D contacts between each 1Mb autosomal bin with the Xist locus (SPRITE data; y axis) and DNA sites enriched by Xist when expressed at high concentration (3x dox; RAP-DNA data; x-axis). **(G)** Relative Xist expression upon RA-induced differentiation (left) and dox-induction (right) of female SHARP-AID mESCs in the absence or presence of auxin as measured by RT-qPCR (primer pair Xist 2). Fold change values were calculated by normalizing to the median of RA differentiated or dox-induced cells in the absence of auxin. Individual points show each replicate value. **(H)** Relative Xist expression in RA differentiated wildtype cells that do not contain dox-inducible Xist (left) and TX cells with dox-induced Xist (right) as measured by RT-qPCR (primer pairs Xist 1, 2). Fold change values were calculated by normalizing to the median of RA differentiated or dox-induced cells in the absence of auxin. Individual points show each replicate value. **(I)** Left: Images of Xist territories in wildtype cells and TX cells; right: % nucleus occupied by Xist in the same cell lines as measured by FISH (Xist) and DAPI (nucleus).

## MATERIALS AND METHODS

### CELL CULTURE, CELL TREATMENTS, AND CELL LINE GENERATION

#### Mouse ES cell culture

Wildtype and endogenous SHARP-HALO-AID TX1072 female mouse embryonic stem cells (mESCs; gift from E. Heard lab) were cultured as previously described [24,39]. Briefly, TX1072 mESCs were grown on gelatin-coated plates in serum-containing ES cell medium (high glucose DMEM (Gibco, Life Technologies), 15% FBS (Omega Scientific), 2mM L-glutamine (Gibco, Life Technologies), 1mM sodium pyruvate (Gibco, Life Technologies), 0.1mM β-mercaptoethanol, 1000 U/ml leukemia inhibitory factor (LIF, Chemicon)), and 2i (3 µM Gsk3 inhibitor CT-99021, 1 µM MEK inhibitor PD0325901). Cell culture medium was replaced every 24 hours.

Expression of Xist and/or each SHARP rescue construct (FL-SHARP, ΔRRM-SHARP, ΔIDR-SHARP, FUS-ΔIDR-SHARP; see Table 1 for complete plasmid list) was induced by treating cells with 2 µg/mL doxycycline (Sigma) for at least 72 hours. Doxycycline-containing medium was replaced every 24 hours. For experiments using SHARP-AID mESCs, cells were treated with indole-3-acetic acid (IAA) for 24 hours before the addition of doxycycline to ensure complete degradation of endogenous SHARP prior to induction of Xist and SHARP rescue constructs. For RNA FISH and immunofluorescence, cells were trypsinized into a single cell suspension, plated directly on poly-D-lysine coated coverslips, and grown for at least six hours before fixation.

#### Human HEK293T cell culture

HEK293T cells were cultured in complete media comprised of DMEM (Gibco, Life Technologies) supplemented with 10% fetal bovine serum (Seradigm Premium Grade HI FBS, VWR), 1X penicillin-streptomycin (Gibco, Life Technologies), 1mM sodium pyruvate (Gibco, Life Technologies), and 1X MEM non-essential amino acids (Gibco, Life Technologies). Cells were maintained in 37oC incubators under 5% CO2.

#### Differentiation with retinoic acid

Wildtype (F1) and TX1072 female mESCs were grown for 24 hours in ES cell medium. ES cell medium was then replaced with MEF medium (high glucose DMEM (Gibco, Life Technologies), 10% FBS (Omega Scientific), 2mM L-glutamine (Gibco, Life Technologies), 1mM sodium pyruvate (Gibco, Life Technologies), 0.1mM MEM non-essential amino acids (Gibco, Life Technologies), 0.1mM β-mercaptoethanol). After 24 hours in MEF medium, the medium was replaced with MEF medium supplemented with 1µM retinoic acid (RA; Sigma). Cells were then grown in MEF medium containing RA for 24 hours (48 hours differentiation total) or 48 hours (72 hours differentiation total). For cells differentiated for 72 hours, MEF medium containing RA was replaced after 24 hours.

To ensure that replacement of the endogenous Xist promoter with a doxycycline-inducible promoter in TX1072 cells does not impair endogenous expression of Xist upon differentiation, Xist levels were measured in both TX1072 and F1 female mESCs (using RT-qPCR; protocol and quantification described below) after 72 hours of differentiation with retinoic acid. Based on this bulk measurement, Xist levels in TX1072 mESCs were approximately half of those in F1 mESCs (Fig. S5H); however, the percent of single cell nuclei occupied by Xist in both TX1072 and F1 mESCs was roughly the same (Fig. S5I; protocol and quantification described below).

#### SHARP-KO cell line generation

To generate a plasmid targeting SHARP for deletion (see Table 1 for complete plasmid list), four different gRNA sequences (see Table 2 for sequences; Fig. S3A) were multiplexed into a Cas9-nickase backbone (Addgene plasmid 48140) as previously described [64]. To create a SHARP knockout (SHARP-KO) cell line, two million TX1072 mESCs were transfected with 1.25 µg of the multiplexed Cas9n-gRNA plasmid containing GFP using the Neon transfection system (settings: 1400 V, 10 ms width, 3 pulses). Successfully transfected cells were enriched by performing FACS for GFP and subsequently plated at low-confluency. After 4-5 days of growth, 96 single colonies were picked and seeded in a 96-well plate. These cells were then split into one plate for PCR genotyping and another plate for maintaining growth until positive clones were identified. PCR genotyping was performed using Q5 High-Fidelity 2X Master Mix (NEB) with the primer pairs listed in Table 2. SHARP-KO clone H8 was used for subsequent experimentation and all other clones were frozen.

#### SHARP rescue lines in SHARP-KO or SHARP-AID parent cells

To generate SHARP rescue cell lines, SHARP rescue constructs (FL-SHARP, ΔRRM-SHARP, ΔIDR-SHARP, FUS-ΔIDR-SHARP; Fig. S3B; see Table 1 for complete plasmid list) were first made using the Gateway LR Clonase system (ThermoFisher). Specifically, ΔRRM-SHARP and ΔIDR-SHARP entry clones were created by modifying a full-length mouse SHARP entry clone using polymerase incomplete primer extension (PIPE) mutagenesis [65]. The specific amino acids deleted in the ΔRRM-SHARP and ΔIDR-SHARP entry clones are as follows.

ΔRRM-SHARP: amino acids 2-590

ΔIDR-SHARP: amino acids 639-3460

These entry clones (FL-SHARP, ΔRRM-SHARP, ΔIDR-SHARP) were then recombined into two different modified versions of the doxycycline inducible PiggyBac destination vector PB-TAG-ERN (Addgene plasmid 80476) containing NGFR (truncated human nerve growth factor receptor) and HALO or eGFP. This destination vector was chosen as it enables stable integration of the rescue constructs by co-transfecting with a PiggyBac transposase [66]. The HALO-tagged version of this plasmid was created by replacing eGFP with NGFR (Addgene plasmid 27489) using Gibson assembly (NEB). HALO was then introduced downstream of rtTA using restriction enzyme digestion and ligation to create PB-HALO-IRES-NGFR. To generate the eGFP-tagged version of this plasmid, HALO was replaced with a 6-HIS-TEV-eGFP sequence using restriction enzyme digestion and ligation. Importantly, eGFP in this construct contains an amino acid substitution (A206K) in order to create a monomeric variant [67].

FUS-ΔIDR-SHARP was generated by recombining the ΔIDR-SHARP entry clone into a modified version of the PB-HALO-IRES-NGFR vector containing the IDR sequence from the FUS protein tagged with mCherry (Addgene plasmid 101223) in place of HALO. Importantly, the IDR from FUS exhibits no sequence homology to endogenous SHARP IDRs (i.e. they have distinct amino acid compositions and distinct proportions of amino acid charge properties), its sequence is ∼10-fold shorter that SHARP IDRs, and the locations of these two IDRs within the SHARP protein are distinct (Fig. S3B).

To generate mESC lines expressing these SHARP rescue constructs, two million SHARP-KO clone H8 or SHARP-AID mESCs were transfected with 2.4 µg of the respective SHARP rescue construct tagged with HALO or eGFP (Table 1), along with 0.8 µg of PiggyBac transposase plasmid (gift from M. Elowitz lab) and 1.2 µg of a non-targeting GFP plasmid (turboGFP; Addgene plasmid 69072 cloned into pcDNA backbone with CMV promoter). Cells that were successfully transfected with the plasmids of interest (SHARP rescue constructs in HALO- or eGFP-tagged PiggyBac destination vector and PiggyBac transposase) were enriched by performing FACS on the co-transfected, non-targeting GFP. Cells were then cultured for 4-5 days to enable the SHARP rescue constructs to stably integrate into the genome (without inducing expression of Xist or the SHARP rescue proteins).

Next, cells were treated with IAA (for SHARP-AID mESCs) and doxycycline (previously described) to induce expression of Xist and the SHARP rescue proteins. Importantly, these cells were cultured in doxycycline for a minimum of 72 hours to ensure that any cells with toxic SHARP expression levels did not survive and were not analyzed further. For HALO-tagged rescue constructs, cells were labeled with 1µM HaloTag Oregon Green Ligand (Promega) according to the manufacturer’s instructions and both the HALO- and eGFP-tagged cell lines were sorted again to enrich for cells expressing the HALO- or eGFP-tagged SHARP rescue constructs (Fig. S3C).

During FACS, laser powers and gains were set based on the lowest expressing samples (FL-SHARP) and these settings were used for all other samples to enrich for cells with comparable expression levels of each rescue construct. Following FACS, cells were kept in medium supplemented with doxycycline and used in further experiments (CLAP, IF, RNA-FISH). Cells were retained only for a maximum of 14 days of culture in doxycycline.

#### Overexpression of SHARP rescue constructs in HEK293T

For those experiments that required high protein expression (live-cell imaging, concentration-dependent imaging assays, CLAP, FRAP), human HEK293T cells were used instead of mESCs because they allow for much higher expression levels and enabled investigation of the biochemical and biophysical properties of each SHARP rescue construct in an independent system that is not undergoing initiation of XCI.

HEK293T cells were transfected using BioT transfection reagent (Bioland) according to manufacturer’s recommendations. Transfected constructs include FL-SHARP, ΔRRM-SHARP, ΔIDR-SHARP, FUS-ΔIDR-SHARP, EED, Ptbp1, or an empty backbone (Table 1); all constructs contained eGFP attached to the N-terminus of each protein of interest driven by a doxycycline-inducible promoter.

For live-cell imaging, fixed imaging, and FRAP (Fig. 2; Fig. S2) ∼10 μg of DNA was used for transfection when cells were grown on a 15cm dish or ∼1 ug of DNA when cells were grown on 3cm glass-bottom dishes (Matek), and DNA concentrations were adjusted to match mole numbers across constructs. 24 hours after transfection, cells were treated with doxycycline (2 µg/mL doxycycline (Sigma)) to induce expression of the proteins of interest and further experiments were performed 48 hours post-doxycycline treatment.

For assays measuring concentration-dependent assembly formation (Fig. 2D, 2E), ∼2.5 fmol of DNA was transfected per well of 24-well plate, adjusting DNA concentration based on the construct being used. 24 hours after transfection, cells were treated with increasing concentrations of dox (0x, 0.1x, 0.5x, 1x where 1x = 2 µg/mL) for 24 hours.

### PROTEIN AND RNA VISUALIZATION

#### Single molecule RNA fluorescence in situ (smRNA-FISH)

RNA-FISH experiments were performed using the ViewRNA ISH Cell Assay (ThermoFisher, QVC0001) protocol with minor modifications. Specifically, cells were fixed on coverslips with 4% formaldehyde in PBS for 15 minutes at room temperature and then permeabilized with 4% formaldehyde and 0.5% Triton X-100 in PBS for 10 minutes at room temperature. Cells were then washed twice with PBS, dehydrated with 70% ethanol, and incubated at -20oC for at least 20 minutes or stored for up to one week. Coverslips were washed twice with PBS and then incubated with the desired combination of RNA FISH probes (Fig. S4A; Table 3; Affymetrix) in Probe Set Diluent at 40oC for at least three hours. Coverslips were then washed once with Wash Buffer, twice with PBS, and once more with Wash Buffer before incubating in PreAmplifier Mix Solution at 40oC for 45 minutes. This step was repeated for the Amplifier Mix Solution and Label Probe Solution. Coverslips were incubated with 1X DAPI in PBS at room temperature for 15 minutes and subsequently mounted onto glass slides using ProLong Gold with DAPI (Invitrogen, P36935).

### Immunofluorescence (IF)

To focus our analysis specifically on nuclear SHARP, pre-extraction was performed on cells prior to immunostaining as previously described [61]. In brief, cells on coverslips were washed once with PBS and then incubated with cold 0.1% Triton X-100 in PBS for 1-3 minutes on ice. Next, cells were fixed on coverslips with 4% formaldehyde in PBS for 15 minutes at room temperature and permeabilized with 0.5% Triton X-100 in PBS for 10 minutes at room temperature. After washing two times with PBS containing 0.05% Tween (PBSt) and blocking with 2% BSA in PBSt for 30 minutes, cells were incubated with primary antibodies overnight at 4oC in 1% BSA in PBSt. After overnight incubation at 4oC, cells were washed 3 times in 1x PBSt and incubated for 1 hour at room temperature with secondary antibodies labeled with Alexa fluorophores (Invitrogen) diluted in 1x PBSt (1:500). Next, coverslips were washed three times in PBSt, rinsed in PBS, rinsed in ddH2O, mounted with ProLong Gold with DAPI (Invitrogen, P36935), and stored at 4oC until acquisition.

Primary antibodies and the dilutions used are as follows: anti-Halo (mouse, Promega G9211, 1:200); anti-Ezh2 (mouse, Cell signaling AC22 3147S, 1:500); anti-SHARP (rabbit, Bethyl A301-119A, 1:200).

### RNA-FISH & Immunofluorescence

For IF combined with in situ RNA visualization, the ViewRNA Cell Plus (Thermo Fisher Scientific, 88-19000-99) kit was used according to the manufacturer’s protocol with minor modifications. First, immunostaining was performed as described above but all incubations were performed in blocking buffer containing RNAse inhibitor from the kit and all wash steps were done in RNAse-free PBS with RNAse inhibitor. Blocking buffer, PBS, and RNAse inhibitors were provided with the kit. After the last wash in PBS, cells underwent post-fixation with 2% formaldehyde in PBS for 10 minutes at room temperature, were washed 3 times in PBS, and then RNA-FISH was performed as described above.

### HALO tag staining

To visualize proteins expressing HALO tags, HaloTag TMR (G8252) or OregonGreen (G2802) was used for fixed sample imaging combined with IF (Fig. S3D) and Janelia549 (GA1110) was used for combined HALO staining and RNA-FISH visualization (Fig. 1B; Fig. S1G). Janelia549 was used for combined HALO staining and RNA-FISH visualization because other HALO ligands did not survive the RNA-FISH protocol. For protein labeling, cells were incubated with HaloTag Ligands according to manufacturer’s instructions and then directly imaged or washed with PBS, fixed in 4% formaldehyde (Pierce, ThermoFisher Scientific), and combined with immunostaining or RNA-FISH.

### IMAGE ACQUISITION AND ANALYSIS

#### Microscopy

Fixed samples were imaged using: Zeiss LSM 800 with the 63x oil objective (RNA-FISH, IF) and collected every 0.3um for 16 Z-stacks, Zeiss LSM 880 with Airyscan with the 63x oil objective (IF) and collected every 0.25um for 20 Z-stacks, or Zeiss LSM 980 with Airyscan2 with the 63x oil objective (IF, RNA-FISH-IF) where zoom, scan format, and number of Z-stacks were optimized based on the software recommendations for the highest resolution (Super-Resolution module). For all images, laser power and gain were set at the beginning of acquisition and remained constant throughout the duration of acquisition to enable comparisons of fluorescent intensities. Live samples were imaged using the Leica Stellaris microscope with 63x water objective (∼80nm xy, ∼300nm z), and 16 Z-stacks were collected every 60 seconds for 5 minutes. The microscope was equipped with a stage incubator to keep cells at 37°C and 5% CO2.

#### Image quantification

Image analysis was performed using ICY or FIJI (ImageJ v2.1.0/1.53c) software. Live-cell movies and 3D reconstructions were created using Imaris software from Bitplane (Oxford Instruments Company).

#### Enrichment over inactive X territory

Xist and SHARP enrichments over the Xist territory (Fig. 1) were quantified using Icy (illustration Fig. S3E). First, a region-of-interest (ROI) was defined that corresponded to the Xist signal across all Z-stacks by applying an intensity threshold (signal above background) and a binary mask was created by demarcating the Xist-coated territory (ROI). Next, several features of these ROIs were quantified, namely: the areas in µm2 (Area), total fluorescent intensities of Xist or SHARP over the entire ROI (Total Intensity), and average fluorescent intensity of Xist or Sharp per area unit of ROI (pixel/interior) (Average Intensity).

SHARP rescue construct enrichments over the Xi demarcated by Ezh2 staining (Fig. 3A, 3B) were quantified using Icy (illustration Fig. S3E). First, images were processed into maximum intensity projections and two types of region-of-interest were specified per each nucleus: i) corresponding to the Xi (X) by creating a binary mask based on Ezh2 marker, ii) and a control region corresponding to a random region (R) of the same size across all Z-stacks. Next, the average fluorescent intensities of SHARP or Ezh2 was quantified per each ROI (X or R). Finally, to normalize for intercellular differences in the expression of rescue constructs, ROI-R was subtracted from ROI-X and divided by ROI-X. As such, if fluorescent intensity signal over the X chromosome (X) is not higher that fluorescent signal in a comparably sized random region in the nucleus (R), the fold change should be centered around 0, whereas when there is enrichment, the signal should be greater than 0.

#### Pattern of SHARP localization

To determine the pattern of SHARP localization after transfecting HEK293T cells with eGFP-SHARP constructs (Fig. 2E, 2G; Fig. S2A), images were first processed into a maximum intensity projection using Icy software. Then, a binary mask was created to demarcate each nucleus of a transfected cell by setting a threshold of eGFP intensity above background levels; all masks were visually verified and, if needed, manually adjusted to fit the nuclear region of cells. Based on these masks, an ROI was defined that corresponded to the entire nucleus. Values for each pixel with the ROI (nucleus) were then extracted and this extracted information was used to quantify total intensity of protein per nucleus (sum of all pixels in an ROI), which corresponds to protein expression levels, and to calculate a SHARP dispersion score describing the differences in the distribution of pixel intensities across the nucleus. Specifically, for each cell, the intensity value at the 99th percentile of the distribution was computed and divided by the mode of the intensity distribution. This score was used because diffused localization shows distributed intensity across the nucleus and non-diffused localization shows accumulation of signal in defined locations, such that the tails of the intensity distributions were much longer. These quantitative assignments were visually confirmed to ensure that these scores capture our definition of diffused and non-diffused organization across cells.

#### Intron RNA FISH

For intron RNA FISH analysis, each image was processed into a maximum intensity projection using Fiji software. Then, the number of spots corresponding to each intron FISH probe per nucleus was manually counted and scored for the presence of Xist signal, number of spots per escape gene (Kdm5c, Kdm6a), and number of spots per silenced gene (Atrx, Pgk1, MeCP2, Gpc4) (Fig. 4A). Because mESCs are known to lose one of the X chromosomes or its fragments while in culture [39,54,55] (Fig. S4B), the analysis was restricted to cells containing two X chromosomes, which were determined by the presence of exactly two spots from escape gene. Additionally, those cells that had more than two spots per any gene or more than one Xist territory per nucleus were excluded from the analysis.

#### Xist percent of nucleus

To calculate the percent of each nucleus occupied by Xist, each image was first processed into a maximum intensity projection using Fiji software. Then, a binary mask was created to demarcate each nucleus by setting a threshold intensity based on DAPI staining; all masks were visually verified and, if needed, manually adjusted to fit the nuclear region of cells. Based on these masks, an ROI was defined that corresponded to the entire nucleus and the size of the nucleus was calculated in FIJI based on the image metadata. Another binary mask was then created to demarcate the Xist territory by setting a threshold intensity based on Xist RNA-FISH staining; all Xist masks were also visually verified and manually adjusted if necessary. An ROI was defined based on these masks and the size of this territory was calculated in FIJI based on the image metadata. The percent of each nucleus occupied by Xist was calculated by dividing the area of the Xist territory by the area of the corresponding DAPI-demarcated nucleus.

#### Fluorescence Recovery After Photobleaching (FRAP)

FRAP experiments were performed in HEK293T cells overexpressing eGFP-tagged FL-SHARP, PTBP1, or EED. 48 hours post-transfection, cells were subjected to FRAP as previously described [68] using the Zeiss LSM 710 with the 40x water objective and equipped with a stage incubator to keep cells at 37°C and 5% CO2. Briefly, in each nucleus ∼1µm2 area was bleached with the Argon laser to quench eGFP and fluorescence recovery was followed while imaging in the GFP channel for 235 seconds. FRAP experiments were analyzed first by measuring the mean fluorescence intensity in the bleached area over time using Icy software and then normalized and averaging over n number of cells (n>5) using EasyFRAP software [69]. Error bars represent standard deviation of at least 5 replicates.

### COVALENT LINKAGE AFFINITY PURIFICATION (CLAP) FOLLOWED BY RNA SEQUENCING

#### Purification of HALO-tagged SHARP

CLAP was performed on mESCs expressing HALO-tagged SHARP constructs (Table 1) as previously described [70] (Fig. 3C, 3D). Briefly, post-transfection, media was removed from cells and then crosslinked on ice using 0.25 J cm−2 (UV2.5k) of UV at 254 nm in a Spectrolinker UV Crosslinker. Cells were collected by scraping in 1X PBS and pelleted by centrifugation. Cell pellets were resuspended in 1 mL of ice cold lysis buffer (50 mM Hepes, pH 7.4, 100 mM NaCl, 1% NP-40, 0.1% SDS, 0.5% Sodium Deoxycholate) supplemented with 1X Protease Inhibitor Cocktail (Promega), 200 U of Ribolock (ThermoFisher), 20 U Turbo DNase (Ambion), and 1X Manganese/Calcium Mix (0.5mM CaCl2, 2.5 mM MnCl2). The samples were incubated on ice for 10 minutes and then at 37°C for 10 minutes at 700 rpm shaking on a Thermomixer (Eppendorf). Lysates were cleared by centrifugation at 15,000 g for 2 minutes, and the supernatant is used for capture. For Halo-protein capture 50 μL of HaloLink Resin was pre-blocked using 1X Blocking Buffer (50 mM HEPES, pH 7.4, 100 μg/mL BSA) for 20 minutes at room temperature with continuous rotation. After incubation, the resin was washed three times with 1X PBSt. The cleared lysate was mixed with 50μl of pre-blocked HaloLink Resin and incubated at 4 °C for 3-16 hours with continuous rotation. The captured protein bound to resin was washed three times with lysis buffer at room temperature and then washed three times at 90°C for 3 minutes while shaking on a Thermomixer at 1200 rpm with each of the following buffers: 1X NLS buffer (1xPBS, 2% NLS, 10 mM EDTA), High Salt Buffer (50 mM HEPES, pH 7.4, 0.1% NP-40, 1M NaCl), 8M Urea Buffer (50 mM HEPES, pH 7.5, 0.1% NP-40, 8 M Urea), Tween buffer (50 mM HEPES, pH 7.4, 0.1% Tween 20) and TEV buffer (50 mM HEPES, pH 7.4, 1 mM EDTA, 0.1% NP-40). Between each wash, samples were centrifuged at 1,000 g for 30 seconds and supernatant was removed. After the last wash, samples were centrifuged at 7,500 g for 30 seconds and supernatant was discarded. For elution, the resin was resuspended in 100 μL of NLS Buffer and 10 μL of Proteinase K (NEB) and the sample was incubated at 50°C for 30 minutes while shaking at 1200 rpm. Capture reactions were transferred to microspin cups (Pierce, ThermoFisher), centrifuged at 2,000 g for 30 seconds, and the elutions were used for RNA purification by RNA Clean and Concentrate-5 kits (Zymo, >17nt protocol).

#### RNA library preparation and sequencing

RNA-seq library preparation was carried out as previously described [71]. Briefly, purified RNA was dephosphorylated (Fast AP) and cyclic phosphates were removed (T4 PNK). The RNA was then cleaned using Silane beads. An RNA adaptor containing a reverse transcription (RT) primer binding site was ligated to the 3’ end of the RNA and the ligated RNA was reverse transcribed into cDNA. The RNA was then degraded using NaOH and a second adaptor was ligated to the single-stranded cDNA. The DNA was amplified, and Illumina sequencing adaptors were added by performing PCR with primers that are complementary to the 3’ and 5’ adapters that were previously added. The molarity of each PCR amplified library was measured using an Agilent Tapestation High Sensitivity DNA screentape and the samples were then pooled at equal molarity. This library pool was then size selected on a 2% agarose gel by cutting between 150-800 nucleotides and performing gel purification (Zymo). To determine the loading density of the final pooled library, the sample was measured using an Agilent Bioanalyzer and Qubit dsDNA High Sensitivity assay (ThermoFisher). The final library was paired-end sequenced on an Illumina HiSeq 2500 with read length 35 × 35 nucleotides.

#### CLAP analysis and visualization

For HALO purifications and RNA binding mapping, sequencing reads were aligned to the mouse genome (RefSeq mm10) using STAR aligner. All low-quality alignments (MAPQ < 255) and PCR duplicates were excluded from the analysis using the Picard MarkDuplicates function (https://broadinstitute.github.io/picard/). The enrichment relative to input coverage across the Xist RNA was quantified by computing the number of reads overlapping the window in the SHARP-elution sample divided by the total number of reads within the SHARP-elution sample. This ratio was normalized by dividing the number of reads in the same window contained in input sample by the total number of reads in the input sample. Because all windows overlapping a gene should have the same expression level in the input sample (which represents RNA expression), the number of reads in the input was estimated as the maximum of either (i) the number of reads over the window or (ii) the median read count over all windows within the gene. This approach provides a conservative estimate of enrichment because it prevents windows from being scored as enriched if the input values over a given window are artificially low, while at the same time accounting for any non-random issues that lead to increases in read counts over a given window (e.g. fragmentation biases or alignment artifacts leading to non-random assignment or pileups). These enrichment values were visualized in IGV [72].

#### Crosslink induced truncation sites

Because UV-crosslinking forms an irreversible covalent crosslink, reverse transcriptase has a well-described tendency to stall at crosslink sites. To exploit this information to identify information about putative protein binding sites at nucleotide resolution, the second adaptor is ligated to the 3’ end of the cDNA. In this way, the start position of the second read in a sequencing pair corresponds to this cDNA truncation point. To quantify these positions, the frequency of reads that start at each nucleotide was counted and plotted along the Xist RNA to identify the positions of direct crosslinking between the protein of interest and the RNA.

### RNA AFFINITY PURIFICATION (RAP) FOLLOWED BY DNA SEQUENCING

#### Cell treatment and preparation

For RAP-DNA sequencing, TX1072 cells were treated with increasing dox concentrations (0.25x, 0.5x, 1x, 2x, 3x where 0x = no dox and 1x = 2 µg/ml) for 72 hours, changing dox containing medium daily. Cells were harvested and crosslinked as previously described [56]. Briefly cells were pelleted, crosslinked with 2 mM disuccinimidyl glutarate for 45 minutes and 3% formaldehyde for 10 minutes, and lysed. Chromatin was then digested to 100-500 bp fragments through a combination of sonication and treatment with TURBO DNase and cell lysates were stored at -80oC until the next step of the procedure.

#### Purification of DNA sites bound by Xist RNA

DNA fragments occupied by Xist RNA were purified for RAP-DNA as previously described [16] with minor modifications. Briefly, the lysate was diluted to hybridization conditions containing 3M guanidine thiocyanate, precleared by adding streptavidin-coated magnetic beads and incubating for 30 minutes at 37°C, mixed with biotin-labeled single stranded DNA capture probes, and incubated at 37°C for 2 hours. 90-mer single stranded DNA oligonucleotide probes spanning the entire length of the target Xist RNA were purchased containing a 5’ biotin (Eurofins Operon) [22]. Next, captured chromatin complexes were eluted with RNaseH and crosslinks were reversed by adding Proteinase K to the probe-bead complexes and incubating overnight at 65°C. Standard Illumina sequencing libraries were generated from eluted DNA fragments and sequenced at a depth of 5-20 million reads per sample of 75-75 or 75-140 long paired-end reads per sample.

#### RAP-DNA analysis and visualization

X to Autosome enrichments were calculated by counting the number of reads that aligned to the X and the number aligned to autosomes (A). This proportion was then compared to the proportion of reads that align to the X or A in the total input sample, which represent the total genomic DNA coverage without any selection. To compute enrichments per region of the genome, the number of reads for each genomic region within 10 kilobase windows was counted and this count was normalized by the total number of sequencing reads within each sample. Each window was then normalized by the proportion measured in the same bin within the input samples. To explore regions on autosomes that contain high Xist coverage, each bin was divided by the median values present on the X chromosome. In this way, all genomic regions containing coverage that was at least as high as half of the regions on the X chromosome could be visualized and their enrichment levels could be directly compared to the overall X chromosome coverage.

#### Computing 3D contact frequencies with the Xist locus

3D contact frequency between individual genomic regions and the Xist transcription locus was calculated as previously described [56]. Specifically, all SPRITE clusters containing a DNA read overlapping the Xist locus (chrX:103460373-103483233, mm10) were extracted and a genome-weight contact frequency was computed by counting the total number of SPRITE clusters for each genomic region within this set. The analysis exclusively focused on clusters containing 2-100 reads per cluster and weighted the contact frequency by the cluster size from which it was present (2/cluster size) as previously described.

#### RT-qPCR

Dox-induced and differentiated female mESCs were lysed in RLT (Qiagen) containing β-mercaptoethanol at a 1:100 dilution. RNA was then isolated using the RNeasy Mini Kit (Qiagen) according to the manufacturer’s instructions. Genomic DNA was removed from the purified RNA samples with TURBO DNase (ThermoFisher) per the manufacturer’s protocol. Total RNA was then purified again using the RNA Clean and Concentrate-5 kit (Zymo, >17nt protocol). cDNA was generated from purified RNA using Maxima H minus reverse transcriptase (ThermoFisher) with random 9mers according to manufacturer’s specifications.

Amplification reactions were run in a Roche LightCycler 480 instrument using LightCycler 480 SYBR Green I Master (Roche) with the primer pairs listed in Table 2. Each sample had between one and six biological replicates and four technical replicates. Median Ct values were used to calculate fold change with the 2-ΔΔCt method. For differentiation and dox induction conditions in the presence or absence of auxin (Fig. 5I), each biological replicate was normalized to the median of the corresponding “with SHARP (-Aux)” condition. For dox-induced samples across increasing concentrations (Fig. S5D), each sample was normalized to the corresponding differentiation (RA) sample. For differentiated wildtype (F1) and TX1072 mESCs (Fig. S5H), each sample was normalized to the corresponding wildtype sample.

#### Other data used in this study

RAP-DNA (F1 2-1 + 48 hours of RA): Xist localization across the X chromosome relative to gene density was measured using our previously published RAP-DNA dataset generated from Xist purification in F1 female mESCs differentiated with retinoic acid for 48 hours [16]. All normalizations and analyses were performed as previously described and plotted using the normalized bedgraphs available at GEO accession GSE46918.

SPRITE (F1 2-1 mESCs): 3D contacts were measured using our previously published RNA-DNA SPRITE dataset [61] that was generated in F1 female mESCs available at GEO accession GSE151515.

#### Data visualization

Bar graphs and violin plots were generated using GraphPad Prism (v8.4.3) or R (v4.0.3). Sequencing data was visualized using IGV (v2.8.11).

#### Statistical tests

To compare distribution of expressed alleles among two populations (Fig. 4) we used two proportions Z-test.

## Notes

### Competing Interest Statement

The authors have declared no competing interest.

